# Variability in β-catenin pulse dynamics in a stochastic cell fate decision in *C. elegans*

**DOI:** 10.1101/245225

**Authors:** Jason R. Kroll, Jasonas Tsiaxiras, Jeroen S. van Zon

## Abstract

During development, cell fate decisions are often highly stochastic, but with the frequency of the different possible fates tightly controlled. To understand how signaling networks control the cell fate frequency of such random decisions, we studied the stochastic decision of the *Caenorhabditis elegans* P3.p cell to either fuse to the hypodermis or assume vulva precursor cell fate. Using time-lapse microscopy to measure the single-cell dynamics of two key inhibitors of cell fusion, the Hox gene LIN-39 and Wnt signaling through the β-catenin BAR-1, we uncovered significant variability in the dynamics of LIN-39 and BAR-1 levels. Most strikingly, we observed that BAR-1 accumulated in a single, 1-4 hour pulse at the time of the P3.p cell fate decision, with strong variability both in pulse slope and time of pulse onset. We found that the time of BAR-1 pulse onset was delayed relative to the time of cell fusion in mutants with low cell fusion frequency, linking BAR-1 pulse timing to cell fate outcome. Overall, a model emerged where animal-to-animal variability in LIN-39 levels and BAR-1 pulse dynamics biases cell fate by modulating their absolute level at the time cell fusion is induced. Our results highlight that timing of cell signaling dynamics, rather than its average level or amplitude, could play an instructive role in determining cell fate.

**Article summary:** We studied the stochastic decision of the *Caenorhabditis elegans* P3.p cell to either fuse to the hypodermis or assume vulva precursor cell fate. We uncovered significant variability in the dynamics of LIN-39/Hox and BAR-1/β-catenin levels, two key inhibitors of cell fusion. Surprisingly, we observed that BAR-1 accumulated in a 1-4 hour pulse at the time of the P3.p cell fate decision, with variable pulse slope and time of pulse onset. Our work suggests a model where animal-to-animal variability in LIN-39 levels and BAR-1 pulse dynamics biases cell fate by modulating their absolute level at the time cell fusion is induced.

## Introduction

During development, cells robustly obtain the correct cell fate to give rise to a viable adult organism, despite internal molecular noise and environmental variability. It is commonly assumed that suppressing this variability is essential for successful development. However, stochastic cell fate decisions, where cells randomly assume one cell fate out of a limited repertoire of different fates, are the cornerstone of many developmental processes (Johnston and Desplan 2010). For example, heterogeneous and stochastic gene expression between cells in the four-cell mouse embryo influences and restricts their cell fates (Goolam et al. 2016). Similarly, photoreceptor cells in the human retina randomly express either a red, green or blue photoreceptor gene (Smallwood, Wang, and Nathans 2002; Roorda and Williams 1999). In these stochastic decision processes, even though each individual outcome is random, the relative frequency of the different cells fates is often tightly controlled.

Currently, stochastic cell fate decisions are best understood in the context of single-celled organisms, where gene expression noise dominates as the key source of variability driving stochastic cell fate decisions (Losick and Desplan 2008; Balaban 2004; Maamar, Raj, and Dubnau 2007; Süel et al. 2006). However, it is unclear how stochastic cell fate decisions are regulated during animal development, as multicellular organisms pose unique constraints compared to single-celled organisms. Here, stochastic cell fate decisions have to be precisely coordinated with developmental timing, are potentially influenced by neighboring cells and rely on external, long-range signals mediated by a small number of key developmental signaling pathways. How these canonical signaling pathways drive stochastic cell fate decisions with strong control over cell fate frequencies is an open question.

Even though *C. elegans* development occurs in a largely invariant manner (Sulston et al. 1983), some cell fate decisions occur in a stochastic manner. One such decision is the specification of the vulval precursor cell (VPC) competence group, beginning early in the L2 larval stage (Myers and Greenwald 2007; Gleason, Korswagen, and Eisenmann 2002). This group consists of six epidermal cells named P3.p-P8.p, which are subsequently patterned to various vulval cell fates by multiple signaling pathways (Gupta, Hanna-Rose, and Sternberg 2012; Hill and Sternberg 1993; D M Eisenmann et al. 1998; Félix 2012; Sternberg and Horvitz 1986; Gleason, Korswagen, and Eisenmann 2002). The establishment of the VPC competence group is partly stochastic, as the P3.p cell assumes VPC fate in roughly 50% of wild-type (N2) hermaphrodites (Fig. 1a), while in the remainder P3.p assumes hypodermal fate by fusing to a neighboring syncytial hypodermal cell, called hyp7 (Gidi Shemer and Podbilewicz 2002; Sternberg and Horvitz 1986). Moreover, the tendency for the P3.p cell to fuse or not in a given strain is sensitive to differences in environmental conditions and genetic backgrounds (Braendle and Félix 2008; Pénigault and Félix 2011a).

**Figure 1.**
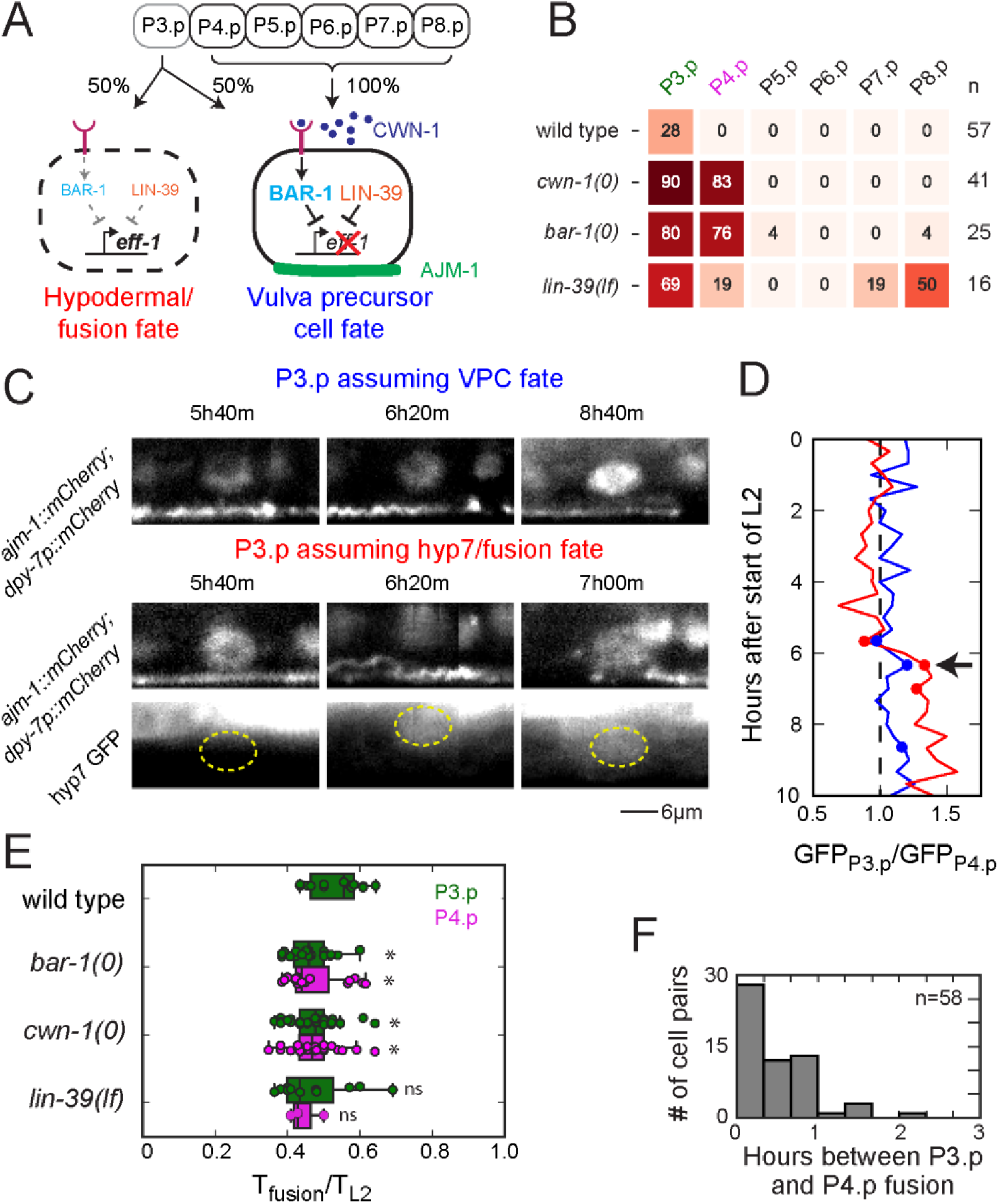
Stochastic cell fate decisions in Pn.p cells. **(A)** Overview of the hyp7/fusion versus vulva precursor cell fate (VPC) decision in the P(3-8).p cells. Cells assuming hyp7/fusion fate fuse (indicated by the dashed line) with the hypodermal syncytium hyp7 and lose the AJM-1 apical junction marker (green). Cell fusion requires the expression of the fusogen EFF-1 and is inhibited by the Hox protein LIN-39 and Wnt signaling through the β-catenin BAR-1. BAR-1 accumulation is induced by binding of Wnt ligands, such as CWN-1 (purple) to Wnt receptors (magenta). **(B)** Measured hyp7/fusion frequencies in Pn.p cells in wild-type and mutant backgrounds. Mutants carried the *ajm-1::GFP* reporter except for *lin-39(lf)* which carried *ajm-1::mCherry*. Wild-type animals carried either *ajm-1:: GFP* (shown here) or *ajm-1::mCherry* (Table 1), with no differences in fusion frequencies. **(C)** AJM-1 dynamics in non-fusing (top) and fusing (bottom) P3.p cells carrying a nuclear *dpy-7p:mCherry* marker. Cell fusion occurred 6h 20m after the start of L2, as shown by the appearance of GFP from the hypodermal syncytium hyp7 in the P3.p nuclear area (region enclosed by yellow line). Simultaneously, AJM-1 showed a pronounced ruffling, followed by its removal from P3.p. In contrast, no such AJM-1 dynamics were observed in non-fusing cells assuming VPC fate. **(D)** Comparing GFP inflow from the hyp7 syncytium in fusing and non-fusing cells as a function of time after the start of the L2 larval stage. Shown is the ratio of GFP fluorescence intensity between P3.p and P4.p in the same animal, where P4.p never fused. The blue and red line corresponds to the non-fusing and fusing cell in (C). Symbols correspond to the time points shown in (C). Arrow indicates the time of AJM-1 ruffling and coincides exactly with inflow of GFP into the fusing cell. **(E)** Individual cell fusion times and box-and-whisker plots for P3.p (green) and P4.p cells (magenta) in different genetic backgrounds. Fusion time was determined for AJM-1 dynamics and is expressed as fraction of the L2 larval stage duration (∼8-12 hrs for all backgrounds). Even though small but significant differences exist in average fusion time between strains, (one-way ANOVA to determine a difference between all strains, followed by Student’s t-test, * indicates P <0.05 in the Student’s t-test), the full distributions show extensive overlap. **(F)** Distribution of difference in cell fusion time between P3.p and P4.p cells, where both cells fuse (data pooled for all genotypes with double fusions).

The Wnt pathway is a highly conserved signaling pathway that regulates many developmental events and cell fates (Ohyama 2006; Hudson et al. 2013; Hirabayashi 2004; Mucenski et al. 2003; Lindström et al. 2014; Clevers and Nusse 2012). Previous investigations into the P3.p cell fate decision showed that its cell fate frequency is extremely sensitive to the dosage of Wnt ligands, particularly *cwn-1*, suggesting that variability in the ligand concentration or in the response of the P3.p cell to Wnt ligands could provide the noise source driving the stochastic fate decision (Pénigault and Félix 2011a; 2011b). In the canonical pathway, the presence of Wnt ligands leads to the accumulation of the transcriptional co-activator BAR-1/β– catenin, which co-regulates Wnt pathway target genes (Sawa and Korswagen 2013; Hendrik C. Korswagen 2002; D M Eisenmann et al. 1998; H C Korswagen, Herman, and Clevers 2000). In addition to the Wnt pathway, mutations in the *C. elegans* Hox gene *lin-39* impact the Pn.p cell fate frequencies, by repression of cell fusion and promoting division of VPC fate cells (Koh et al. 2002; Gidi Shemer and Podbilewicz 2002; Maloof and Kenyon 1998; Roiz et al. 2016; Clark, Chisholm, and Horvitz 1993). Both Wnt signaling and LIN-39 inhibit hyp7/fusion fate, with loss-of-function mutants exhibiting increased frequency of cell fusion, including in the P4.p-P8.p cells that otherwise never assume hyp7/fusion fate (Gleason, Szyleyko, and Eisenmann 2006; Myers and Greenwald 2007). However, what aspects of Wnt signaling and LIN-39 dynamics control the frequency of hyp7/fusion versus VPC fate in P3.p remains unknown.

Here, we use a novel time-lapse microscopy approach (Gritti et al. 2016) to observe gene expression and signaling dynamics in single Pn.p cells during specification of the VPCs, allowing us to directly connect variability during the decision process to the final cell fate outcome. Using this approach, we observed animal-to-animal variability in LIN-39 and BAR-1 dynamics that could be linked to cell fate outcome. In particular, we found that BAR-1 accumulated in a single, ∼1-4 hr pulse in the Pn.p cells at the time of the hyp7/fusion versus VPC fate decision, with strong variability in pulse slope and timing. Using experiments in mutants combined with quantitative data analysis, we arrived at a model where stochastic variability in LIN-39 levels and BAR-1 pulse dynamics biases cell fate by modulating their absolute level at the time cell fusion is induced.

## Results

### Time-lapse microscopy of a stochastic cell fate decision

So far, whether P3.p undergoes fusion or assumes VPC fate in wild-type or mutant animals has been assessed only after the process has completed (Myers and Greenwald 2007; D M Eisenmann et al. 1998; S Alper and Kenyon 2001; Scott Alper and Kenyon 2002; Pénigault and Félix 2011b; 2011a; Chen and Han 2001). To correlate early stochastic molecular events to eventual cell fate outcome it is essential to follow these processes directly in time. Here, we utilize a fluorescent time-lapse microscopy approach we developed recently to study single-cell dynamics inside moving and feeding *C. elegans* larvae for their entire ∼40 hr development (Gritti et al. 2016). We tested whether we could directly observe P3.p fusion events in single animals. We initially used two measures of cell fusion: first, imaging the apical junction protein AJM-1, which localizes on the apical junction of Pn.p cells but is displaced upon cell fusion and has been used as a cell fusion marker in other studies, including live imaging applications (Brabin, Appleford, and Woollard 2011; G Shemer et al. 2004). Using animals carrying AJM-1::GFP, we found that the fusion event occurred in 28% of animals during the L2 stage (n= 57 total animals, see Table 1), and could detect disruption of AJM-1 and its subsequent loss, indicating that the P3.p fusion fate is indeed stochastic between animals in the same strain. The *ajm-1::mCherry* strain showed similar results (Table 1). In both strains, the cell fusion fate of the P3.p cell was reduced from the canonical 50% ratio, possibly resulting from the fluorescent AJM-1 markers, or the environmental conditions of our time-lapse microscope (Braendle and Félix 2008). Second, as an independent measure of cell fusion, we directly observed the flow of GFP from the hypodermis into the fused P3.p cell, using animals carrying an extrachromosal array targeting GFP expression to the hyp7 hypodermal syncytium, while simultaneously monitoring the apical junction marker AJM-1::mCherry. As seen before, the AJM-1::mCherry signal expanded along the A-P axis during the early L2 larval stage (Supplementary Movies S1-S2, Fig. 1c). In animals with a fusing cell, this was followed by a sudden and pronounced ruffling of the AJM-1:mCherry signal and a rapid retraction of AJM-1::mCherry towards the posterior, with its fluorescent signal fully disappearing from P3.p within 1 hr (Fig. 1c). Inflow of GFP from the hypodermis into P3.p was observed as soon as AJM-1::mCherry retraction commenced (Fig. 1c,d), showing that both are accurate markers of (the time of) fusion. Because AJM-1 fluorescence was easily monitored in combination with other quantification, we used AJM-1 dynamics to establish fate and timing of P3.p fusion for all subsequent experiments.

**Table 1.**

| Genotype | Fusion/hyp7cell fate rates (%) <sup>a</sup> |  |  |  |  |  |  |
| --- | --- | --- | --- | --- | --- | --- | --- |
|  | P3.p | P4.p | P5.p | P6.p | P7.p | P8.p | N |
| *ncls13[ajm-1::GFP]* <sup>b</sup> | 28 | 0 | 0 | 0 | 0 | 0 | 57 |
| *ouls20[ajm-1::mCherry]* <sup>b</sup> | 37 | 0 | 0 | 0 | 0 | 0 | 30 |
| *cwn-1(ok546);ncls13[ajm-1::GFP]*<br>***cwn-1(0)*** | 90 | 83 | 0 | 0 | 0 | 0 | 41 |
| *bar-1(ga80);ncls13[ajm-1::GFP]*<br>***bar-1(0)*** | 80 | 76 | 4 | 0 | 0 | 4 | 25 |
| *lin-39(gk893);ncls13[ajm-1::GFP]* <sup>c</sup><br>***lin-39(0)*** | 100 | 100 | 100 | 100 | 100 | 100 | 17 |
| *lin-39(n709);ouls20[ajm-1::mCherry]*<br>***lin-39(lf)*** | 69 | 19 | 0 | 0 | 19 | 50 | 16 |
| *lin-39::GFP;HIS24-H2B::mCherry; ncls13[ajm-1::GFP]*<br>***lin-39(++)*** | 2 | 0 | 0 | 0 | 0 | 0 | 48 |
| *cwn-1(ok546);lin-39::GFP;HIS24-H2B::mCherry;ncls13[ajm-1::GFP]*<br>***cwn-1(0); lin-39(++)*** | 20 | 14 | 1 | 0 | 0 | 0 | 126 |
| *bar-1::GFP;ouls20[ajm-1::mCherry]*<br>***bar-1(++)*** | 0 | 0 | 0 | 0 | 0 | 0 | 30 |
| *bar-1::GFP;cwn-1(ok546);ouls20[ajm-1::mCherry]*<br>***cwn-1(0); bar-1(++)*** | 6 | 8 | 0 | 0 | 0 | 0 | 64 |
| *bar-1::GFP;lin-39(n709);ouls20 [ajm-1::mCherry]*<br>***lin-39(lf); bar-1(++)*** | 24 | 4 | 3 | 0 | 13 | 19 | 70 |
| *bar-1::GFP<sup>CRISPR</sup>; ouls20[ajm-1::mCherry]*<br>***bar-1::GFP<sup>CRISPR</sup> <sup>b</sup>*** | 30 | 0 | 0 | 0 | 0 | 0 | 23 |

Pn.p fusion frequencies in different genetic backgrounds with time-lapse microscopy techniques.
<sup>a</sup> Fusion rates are rounded to the nearest percentage. Fusion events were counted by the loss of *ajm-1* staining during the L2 stage, and non-fusion animals were only counted if the animal reached the L3 ecdysis without a fusion event.
<sup>b</sup> No statistical difference between P3.p fusion rates in these control strains, (Chi-square statistic =0.68, P= 0.71).
<sup>c</sup> P3.p – P8.p fused prematurely during the L1 stage in the null mutant.

Even though changes to the frequency of P3.p hyp7/fusion versus VPC fate in mutants are well studied (Pénigault and Félix 2011b; 2011a; Chen and Han 2001; Myers and Greenwald 2007; D M Eisenmann et al. 1998; S Alper and Kenyon 2001; Scott Alper and Kenyon 2002), it was not known how such mutants impact the timing of this decision. We quantified the time of P3.p fusion in wild-type animals and found that cell fusion occurred in a relatively narrow time window between 40-60% of the L2 larval stage (Fig. 1e). We then examined mutants in which fusion frequency is increased by removing inhibitory Wnt signaling or LIN-39 (Fig. 1b). We found that in these mutants P3.p fusion occurred within the same time window as wild-type animals, with only small differences between wild-type and mutant animals in average timing (Fig. 1e). Strikingly, even though the exact time of fusion can vary as much as 2 hrs between animals, when multiple VPCs fused in a single animal, they typically did at the same time (Fig. 1f). In the absence of key repressors of hyp7/fusion fate, cell fusion frequency is increased independently of its timing, providing evidence that hyp7/fusion inhibitors (Wnt signaling, LIN-39) do not control timing of fusion, but rather modulate hyp7/fusion frequency. This also suggests that a yet-unknown signal exists that activates cell fusion at the appropriate time.

### Stochastic *eff-1* induction precedes P3.p fusion

The most downstream regulator of the hyp7/fusion versus VPC fate decision is the gene *eff-1*, a fusogenic protein whose expression is sufficient to induce cell fusion (Mohler et al. 2002; G Shemer et al. 2004). EFF-1 is a transmembrane protein that is required for most cell fusions in *C. elegans*, and must be present on both the Pn.p and hyp7 plasma membrane to induce fusion (Zeev-Ben-Mordehai et al. 2014; Smurova and Podbilewicz 2016). To understand if cell fate frequency is regulated on the level of *eff-1* expression, and how it might be regulated by the Hox/LIN-39 and Wnt signaling pathways, we counted *eff-1* mRNA molecules in Pn.p cells,using single molecule FISH (smFISH) (Raj et al. 2008). In wild-type animals in the late L1 and early L2 stage (230 - 325 μm in body length), stages of development immediately preceding the hyp7/fusion versus VPC fate decision, we found that all P3.p cells were unfused, as determined by the presence of the AJM-1 signal. They also exhibited low *eff-1* expression in P3.p, <10 molecules (Fig. 2a,d), similar to the P4.p cell that always assumes VPC fate in wild-type animals. However, in older, mid-L2 stage animals (>325 μm in length), corresponding to the time of fusion, we observed a subset of animals expressing ∼30-50 molecules in unfused P3.p cells, something not observed in P4.p (Fig. 2b,d). In fused P3.p cells, we found similar number of *eff-1* mRNA molecules located in close proximity to the cell nucleus, suggesting that high *eff-1* expression is maintained after fusion, before finally disappearing by the end of the L2 stage (Supplemental Fig. S1a,b). We confirmed that high *eff-1* expression preceded cell fusion by examining a temperature-sensitive loss-of-function point mutation in *eff-1* (Mohler et al. 2002). Here, we still found high *eff-1* mRNA levels in P3.p at the restrictive temperature, even though cell fusion was blocked (Supplemental Fig. S1c).

**Figure 2.**
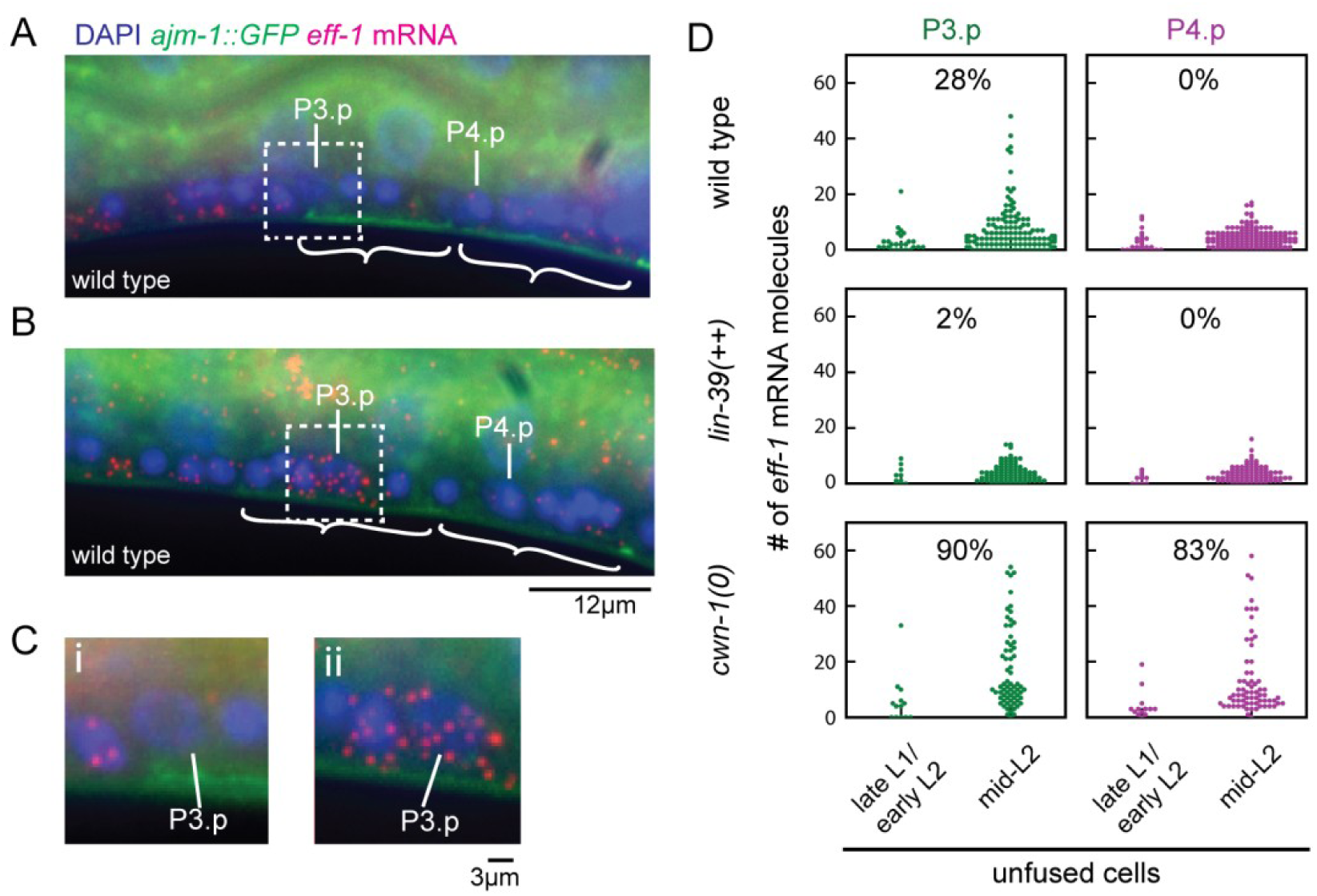
Stochastic *eff-1* expression in Pn.p cells precedes cell fusion. **(A)**,**(B)** Example of (A) low and (B) high *eff-1* expression preceding cell fusion in wild-type animals of similar age, as determined by body length. Single *eff-1* mRNA molecules (red) were visualized using smFISH and nuclei with DAPI (blue). Unfused cells were selected based on AJM-1::GFP localization (green), with white brackets indicating intact apical junctions typical of unfused cells. **(C)** Panels (i) and (ii) show details of the P3.p cell region, corresponding to the area delineated by the dashed line in panels (A) and (B). **D)** Number of *eff-1* mRNA molecules in unfused P3.p and P4.p cells, for wild type animals (n=184) and mutants with decreased (*lin-39(*++*), n=99*) and increased (*cwn-1(0), n=81*) cell fusion frequency. Late L1 and early L2 animals (230-325 μm body length) are at a developmental stage preceding the hyp7/fusion versus VPC fate decision, with fusion occurring only in mid-late L2 animals (>325 μm body length). In each panel, the fusion rate of that cell in the indicated genotype is indicated as a percentage.

The fraction of wild-type animals showing high (>20 molecules) *eff-1* mRNA levels in P3.p was significantly smaller than the expected fraction of animals where P3.p assumes hyp7/fusion fate. For our analysis, we randomly sampled animals within the time window we expected fusion to occur. Given the observed variability in the time of fusion (Fig. 1f), it is expected that some animals with low *eff-1* expression would have ultimately fused at a later point. In particular, the fraction of animals observed with high *eff-1* expression in unfused P3.p cells should increase with the average duration the cell expresses high *eff-1* before this results in fusion. Hence, our results suggested that induction of high *eff-1* expression was quickly followed by cell fusion.

To understand how *eff-1* expression is correlated to cell fate frequency, we quantified *eff-1* levels in a strain with a functional LIN-39::GFP insertion that caused low fusion frequency (*lin-39(*++), ∼2% P3.p fusion frequency). Consistently, we no longer observed unfused P3.p cells expressing high (>20 molecules) *eff-1* levels (Fig. 2d). In contrast, in the *cwn-1(0)* mutant that lacks the dominant Wnt ligand and exhibited high (>80%) fusion frequency both in P3.p and P4.p, we found that the numbers of unfused cells observed with high *eff-1* expression had increased substantially, most strikingly in P4.p (Fig. 2d). Together, this indicates that Wnt signaling and LIN-39 controlled cell fate frequency mainly by tuning the fraction of cells in which high *eff-1* expression is induced.

### Bias of cell fate decision by noise in LIN-39 protein level

The Hox transcription factor LIN-39 inhibits hyp7/fusion fate by repressing *eff-1* expression (Gidi Shemer and Podbilewicz 2002), with *lin-39* null mutations causing all Pn.p cells to fuse in the L1 larval stage (Clark, Chisholm, and Horvitz 1993; Wang et al. 1993). Hence, stochastic variability in LIN-39 protein levels could result in variability in induction of high *eff-1* expression. It was shown previously that LIN-39 levels are similar between P3.p and P4.p in early L2 larval stage animals prior to cell fusion (Pénigault and Félix 2011a), even though both cells have a different fate frequency (Fig. 1b). However, LIN-39 dynamics in individual cells were not followed over time and linked to the eventual cell fate. Hence, it remains possible that small differences in LIN-39 between P3.p cells in different animals are sufficient to explain the outcome. To connect animal-to-animal variability in LIN-39 level with P3.p cell fate, we performed time-lapse microscopy on animals carrying an integrated low-copy *lin-39::GFP* translational fusion transgene (Sarov et al. 2012) and *ajm-1::GFP* as a cell fusion marker (Fig. 3a,b). Since *lin-39::GFP (lin-39(*++)) is present as an insertion, it decreased the P3.p fusion rate from ∼30% to ∼2%, making it challenging to capture sufficient P3.p fusion events for analysis. For that reason, we further crossed this reporter into the *cwn-1(0)* mutant, which increased the P3.p and P4.p fusion rates to 20% and 14%, respectively (Table 1).

**Figure 3.**
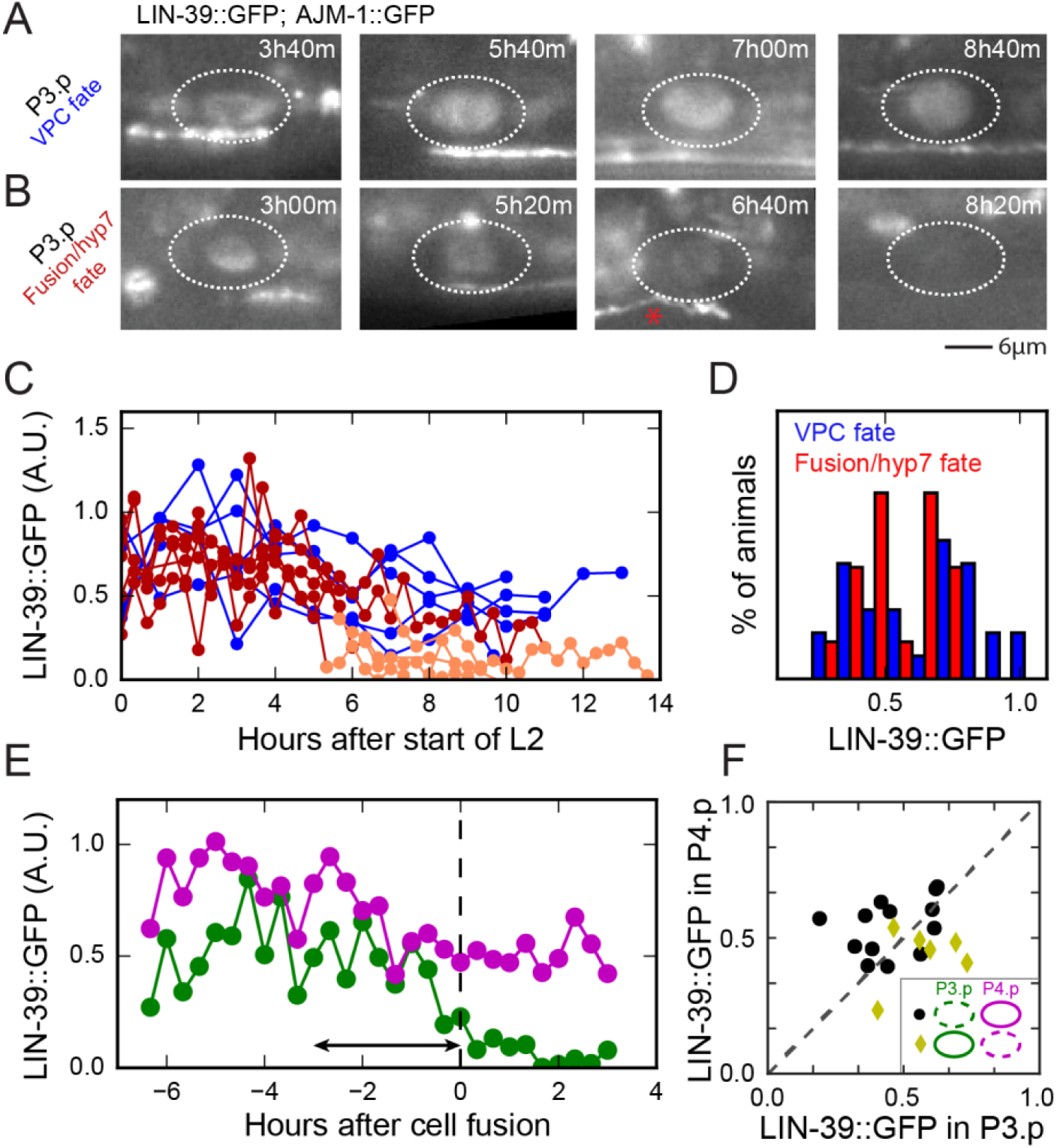
Variability in LIN-39 protein level. **(A), (B)** Image sequence of a cell assuming (A) VPC or (B) hyp7/fusion fate. Nuclear LIN-39 levels are visualized using a *LIN-39::GFP* protein fusion, while cell fusion is determined using an *ajm-1::GFP* reporter. The red star indicates cell fusion has occurred. Times are relative to the start of the L2 stage. **(C)** Nuclear LIN-39 fluorescence in fusing (red) and non-fusing (blue) P3.p cells. For fusing P3.p cells, LIN-39 levels are variable prior to fusion (dark red) and fall rapidly after fusion (light red). **(D)** Distribution of nuclear LIN-39 fluorescence in fusing (red) and non-fusing (blue) P3.p cells show strong overlap. In fusing cells, fluorescence was averaged over 3 hour window directly prior to fusion, whereas in non-fusing cells a time window of 3 hours prior to the average time of cell fusion was used (N = 48 total animals) **(E)** Nuclear LIN-39 fluorescence in P3.p (green) and P4.p (magenta) in an animal in which P3.p but not P4.p assumes hyp7/fusion fate. The arrow indicates the time window over which LIN-39 fluorescence is averaged in (D) and (F). The dashed line indicates the time of fusion. (F) Nuclear LIN-39 fluorescence levels in P3.p and P4.p for animals in which P3.p assumes hyp7/fusion fate and the P4.p assumes VPC fate (black circle) or the reverse (yellow diamond), see legend for schematic of P3.p and P4.p fates, dashed circle indicates fusion. Each symbol corresponds to a single animal and is averaged over a 3 h time window prior to cell fusion. In general, LIN-39::GFP levels were lower in the cell that assumed hyp7/fusion fate.

We observed LIN-39 in the P3.p nucleus at the start of the L2 larval stage and it remained there for the entire larval stage when P3.p assumed VPC fate (Fig. 3a,c). However, in P3.p cells that fused, nuclear LIN-39 levels decreased rapidly after fusion commenced and fully disappeared within 90 mins (Fig.3b,c), consistent with past observations of loss of LIN-39 in fused Pn.p cells (Pénigault and Félix 2011a). LIN-39 levels were not significantly different in the *cwn-1(0)* mutant compared to wild type, indicating that the Wnt ligand acts in a different pathway from LIN-39 to inhibit *eff-1* expression. We compared the distribution of LIN-39 levels, averaged over 3 hrs prior to fusion in P3.p cells that assumed hyp7/fusion fate, with the distribution in P3.p cells that assumed VPC fate, averaged over 3 hrs prior to the average time of P3.p fusion in this strain (Fig. 3d). We found strong overlap between the two distributions, also when changing the size of the time window (data not shown), making it unlikely that fluctuations in LIN-39 levels are the main driver of *eff-1* induction and cell fusion.

These results leave open the question whether the observed variability in LIN-39 has any effect on the P3.p cell fate outcome. In *lin-39::GFP; cwn-1(0)* animals, P4.p also assumed hyp7/fusion fate in a stochastic manner, allowing us to test whether differences in LIN-39 levels between P3.p and P4.p within the same animal are correlated with their eventual fate. First, we established that in this strain LIN-39 distributions for P3.p and P4.p were similar and also showed substantial overlap between fusing and non-fusing cells. We then selected animals in which one cell, either P3.p or P4.p, fused but the other assumed VPC fate (Fig. 3e). Indeed, in these animals, absolute LIN-39 level was not predictive of the eventual fate of P3.p or P4.p, but the difference in LIN-39 levels between P3.p and P4.p was predictive of their fate (Fig. 3f), with fusing cells having lower LIN-39 levels than their non-fusing neighbor cell (P = 0.0095, Fisher’s Exact Test). This shows that variability in LIN-39 levels is correlated to the hyp7/fusion versus VPC fate decision, but likely in conjunction with another source of variability.

### β-catenin activation dynamics during the cell fate decision

Upon activation of Wnt receptors by Wnt ligands, β-catenin accumulates in the cell and nucleus (Sawa and Korswagen 2013). In the absence of Wnt ligands, β-catenin is degraded by a degradation complex. In P3.p, the presence of the β-catenin BAR-1 is required to inhibit *eff-1* expression and, hence, inhibit hyp7/fusion fate (D M Eisenmann et al. 1998). In contrast to β– catenins involved in the Wnt asymmetry pathway (Mila et al. 2015; Park and Priess 2003), dynamics of BAR-1 during canonical Wnt signaling is poorly characterized.

We aimed to quantify activation of the Wnt pathway by monitoring the accumulation dynamics of BAR-1/β-catenin. We first endogenously tagged the C-terminus of BAR-1 with GFP using CRISPR/Cas9 genome editing, (see Supplemental Methods) (Vicencio et al. 2019). The fusion rate when the strain was crossed to the *ajm-1::mCherry* marker was similar to other controls, (see Table 1), indicating that tagging BAR-1 with GFP has no detrimental effect on its function. However, no fluorescence was detected in the Pn.p cells with the laser intensities and exposure times required for our time-lapse method. To visualize BAR-1 dynamics, we instead used a multi-copy integrated BAR-1::GFP transgene, *gaIs45*, which rescues the *bar-1(0)* phenotype (D M Eisenmann et al. 1998) and has been used previously to study BAR-1 dynamics during male hook development (Yu et al. 2009). Similar transgenes have been used extensively to study the dynamics of the *C. elegans* β-catenins SYS-1 (Robertson et al. 2017) and WRM-1 (Kim et al. 2013). A disadvantage of using a multi-copy insertion is that the elevated BAR-1 level perturbs the observed P3.p fate frequency. However, a key advantage is its increased fluorescence signal, which was crucial for imaging its dynamics using our time-lapse imaging approach.

Wnt ligand gene expression levels and protein distribution are considered to be largely constant in time (Pani and Goldstein 2018; Coudreuse et al. 2006). Hence, we expected BAR-1 to show constant expression and dynamics in P(3-8).p cells, similar to LIN-39. Instead, we found that BAR-1::GFP levels were strikingly dynamic, with no BAR-1::GFP in P(3-8).p at the start of the L2 stage, followed by steady accumulation of BAR-1::GFP in P(3-8).p at the mid-L2 stage that lasted 1-4 hours and was strongly coordinated between cells (Supplemental Movie S3, Fig. 4a-c). After the accumulation phase, BAR-1 was rapidly degraded, with the overall dynamics of BAR-1 resembling a single pulse. The protein was detected in both the nucleus and cytoplasm. We found stochastic variability in the amplitude of the BAR-1 pulse between different Pn.p cells (Fig. 4b,c). It was speculated that P3.p, which is considered most distant to the source of Wnt ligands, receives a lower Wnt signal than P(4-8).p, thereby resulting in its occasional hyp7/fusion fate (Harterink et al. 2011; Pénigault and Félix 2011b). However, the BAR-1::GFP accumulation pulse in P3.p was frequently of similar or higher amplitude compared to the other Pn.p cells and, in general, we found no sign of a systematic spatial pattern in downstream Wnt signaling. We also found significant variability in the amplitude and timing of the BAR-1::GFP pulse in the P3.p cell compared between different animals (Fig. 4d).

**Figure 4.**
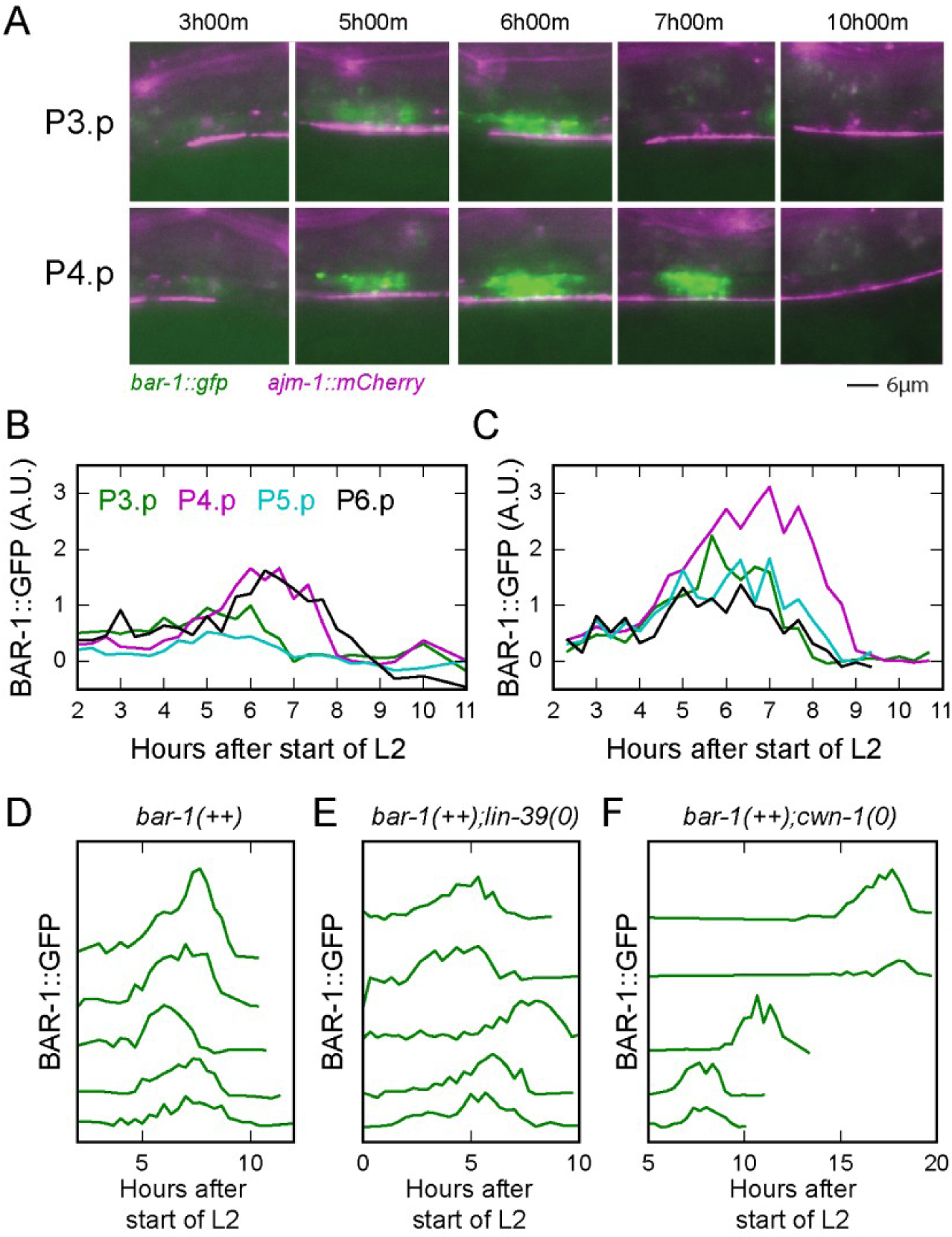
BAR-1 dynamics during cell fate decision. **(A)** Image sequence of BAR-1::GFP dynamics (green) over time in P3.p and P4.p cells of a *bar-1*(++); *ajm-1::mCherry* animal, with both cells assuming VPC fate. **(B)**,**(C)** Dynamics of cellular BAR-1::GFP fluorescence in P(3-6).p cells in two different animals. The animal in (B) corresponds to images in (A). Different cells show similar timing of BAR-1 pulse onset, but vary in amplitude. **(D)-(F)** Examples of BAR-1::GFP dynamics in non-fusing P3.p cells in (D) *bar-1(*++), (E) *bar-1(*++*);lin-39(lf)*, (F) *bar-1(*++*);cwn-1(0)* animals. Traces for different animals are shifted along the vertical axis for clarity. Pulses show strong differences in time of onset and duration, even within the same strain. Traces run until the end of the L2 larval stage. Some *bar-1(*++*);cwn-1(0)* animals displayed significantly extended L2 stage duration (∼15-20hr). Total animals imaged for *bar-1(*++*), bar-1(*++*);lin-39(lf)*, and *bar-1(*++*);cwn-1(0)* genotypes are n= 30, 64, and 70 animals, respectively.

Since the BAR-1::GFP reporter is integrated as a functional multi-copy transgene, we expected this strain to act as a BAR-1 overexpression mutant and, indeed, observed no P(3-8).p cell fusions (n=30 animals, Table 1). Hence, we refer to this strain as *bar-1(*++). To study how BAR-1 dynamics relates to cell fate frequency, we used different approaches to increase the frequency of hyp7/fusion fate. First, we decreased the level of the inhibitor LIN-39, using the *lin-39(n709)* temperature sensitive loss-of-function mutant (Table 1). We found that *bar-1(*++*);lin-39(lf)* animals showed similar BAR-1::GFP dynamics (Fig. 4e). Next, we sought to lower BAR-1 levels in the *bar-1(*++) background by decreasing activity of the Wnt signaling pathway, using the *cwn-1(0)* mutant that lacks the CWN-1 Wnt ligand (Fig. 1a). In these animals, the BAR-1::GFP pulse amplitude was lower (Fig. 4f), and in some animals the L2 stage was significantly lengthened. Moreover, we found that the BAR-1::GFP pulse occurred at significantly later times, as a fraction of larval stage duration, even in animals with a L2 larval stage duration similar to wild-type. The BAR-1 pulse also showed considerable variability in timing and amplitude in these mutants. Finally, in both strains, fusing Pn.p cells did so during the early accumulation phase of the BAR-1 pulse (Fig. 5a, n=29 cells). Even though the increased BAR-1 protein levels in the *bar-1(*++*);lin-39(lf)* and *bar-1(*++*);cwn-1(0)* mutants do not represent the wild-type condition, these animals still undergo a stochastic hyp7/fusion fate decision, allowing us to correlate pulse dynamics with cell fate outcomes.

**Figure 5.**
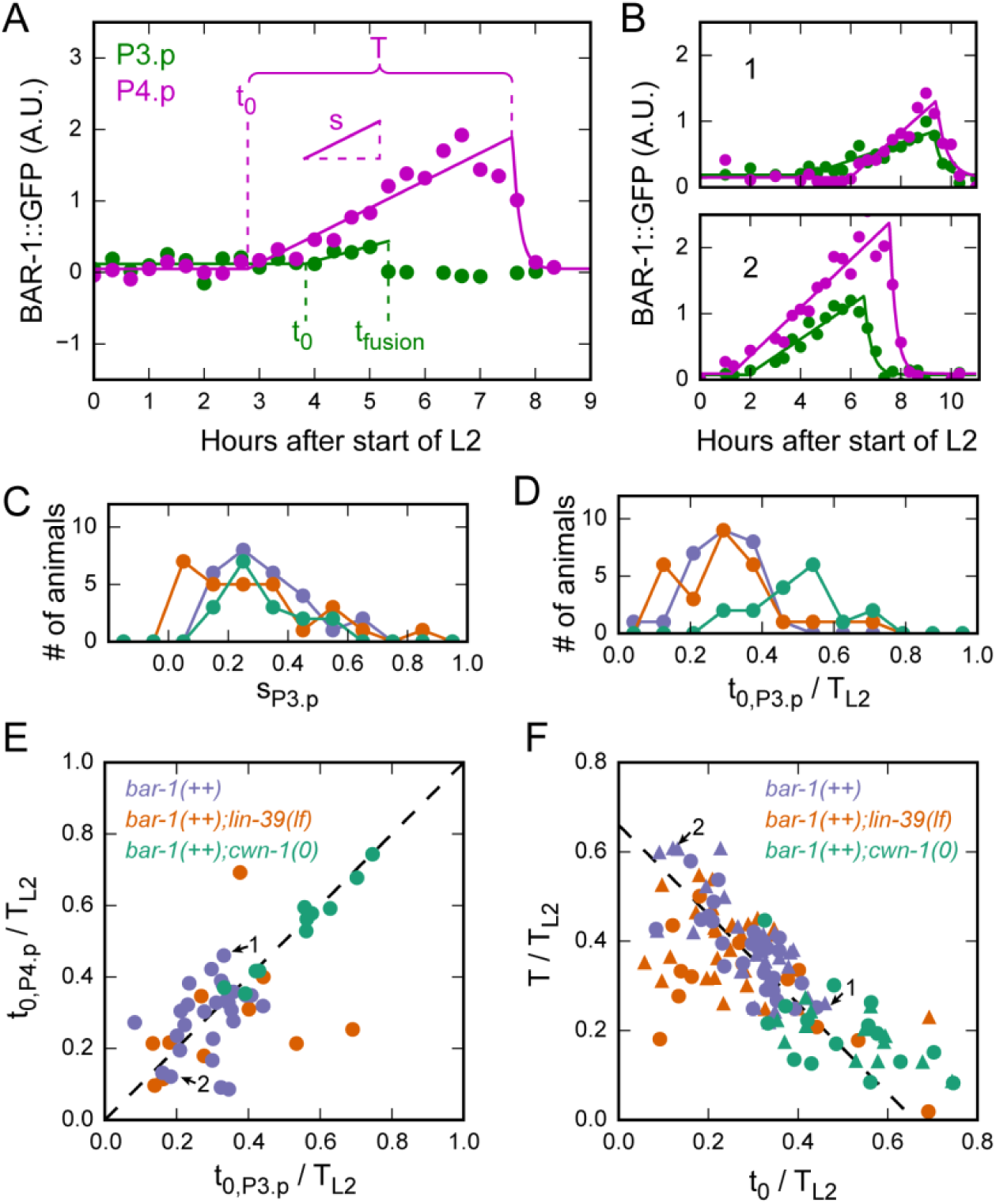
Variability in BAR-1 dynamics. **(A)** Quantifying BAR-1 pulse dynamics in a *bar-1(*++*);lin-39(lf)* animal. Symbols are measured BAR-1::GFP levels in P3.p (green) and P4.p (magenta). Here, P3.p, but not P4.p, assumed hyp7/fusion fate. Solid lines represent a model of BAR-1 accumulation dynamics (Eq. 3 in Methods). Using the model, each BAR-1 pulse is described by three key parameters: pulse onset time *t*_0_, pulse slope and pulse duration *T* (for non-fusing VPCs) or time of fusion *t*_fusion_ (for hyp7/fusion cells). **(B)** Animals showing differences in (relative) timing of BAR-1 accumulation pulses. **(C), (D)** Distribution of (C) pulse slope and (D) pulse onset time *t*_0_ in P3.p cells, in *bar-1(*++) (purple, n= 27 animals), *bar-1(*++*);lin-39(lf)* (orange, n= 30 animals) and *bar-1(*++*);cwn-1(0)* mutants (green, n= 17 animals). *T*_L2_ is the L2 larval stage duration. Strains differ in BAR-1 pulse onset time rather than pulse slope. **(E)** Correlation in pulse onset time between P3.p and P4.p cells. Each symbol corresponds to a P3.p and P4.p cell in a single animal where both assumed VPC fate. Arrows indicate the animals in (B). **(F)** Correlation between pulse onset time *t*_0_ and pulse duration *T* in non-fusing P3.p (circles) and P4.p cells (triangles). The dashed line is *T/T*_L2_ = 0.66 − *t*_0_*/T*_L2_ BAR-1 pulses that start later in the L2 larval stage have a shorter duration.

### Variability in β-catenin dynamics

To characterize variability in BAR-1 accumulation dynamics, we used a minimal parameterization of the BAR-1 pulse shape to fit to the experimental data (Fig. 5a, Eq. 3 in Methods). Briefly, we assume that prior to the BAR-1 pulse, Wnt signaling is inactivated and BAR-1 is degraded. At pulse onset time *t*_0_, Wnt signaling is activated, leading to inhibition of BAR-1 degradation and hence linear accumulation of BAR-1 in the cell. Linear BAR-1 accumulation continues for a pulse duration *T* in cells that assume VPC fate or until the time of fusion, *t*_fusion_, in cells that assume hyp7/fusion fate. Upon fusion BAR-1 vanishes immediately, as observed experimentally, whereas in cells that assume VPC fate, BAR-1 levels decrease exponentially once the pulse ends. This fitted the experimental data surprisingly well (Fig. 5a,b). Moreover, it allowed us to describe each BAR-1 pulse by three parameters: pulse onset time *t*_0_, pulse slope *s* and pulse duration *T* for VPC cells or fusion time *t*_fusion_ for hyp7/fusion cells.

First, we compared the distribution of pulse onset time *t*_0_ and linear slope of each BAR-1 accumulation pulse between strains. We found that both were highly variable among animals (Fig. 5c,d). We found that the pulse parameters of *bar-1(*++) and *bar-1(*++*);lin-39(lf)* were similar, consistent with increase in hyp7/fusion fate frequency in this mutant resulting from the absence of the fusion inhibitor LIN-39 rather than changes in Wnt signaling. In contrast, we assumed that the increase in frequency of hyp7/fusion fate in *bar-1(*++*);cwn-1(0)* animals was due to a decrease in BAR-1 level, given that BAR-1 accumulation is thought to be proportional to the amount of external Wnt ligands. Surprisingly, we found that the pulse slope distribution was highly similar for *bar-1(*++) and *bar-1(*++*);cwn-1(0)* animals (Fig. 5c) and that the only different characteristic was the delayed pulse onset (Fig. 5d).

In general, we found that pulse onset time *t*_0_ was correlated between P3.p and P4.p in the same animal, meaning that if BAR-1 accumulation started late in the L2 larval stage in P3.p, it was also likely to start late in P4.p (Fig. 5e). However, in *bar-1(*++) and *bar-1(*++*);lin-39(lf)* animals we still observed significant variability in pulse onset time between P3.p and P4.p in the same animal, with BAR-1 accumulation in P3.p preceding that in P4.p as often as the reverse (Fig. 5e). Strikingly, in the *bar-1(*++*);cwn-1(0)* mutant the variability in pulse onset time between P3.p and P4.p was almost completely removed, with the onset of BAR-1 accumulation occurring in P3.p and P4.p within 20 min in all animals (Fig. 5e). This result suggests that the Wnt ligand *cwn-1* not only controls the average onset of BAR-1 pulses, but it also induces variability in pulse onset time between P3.p and P4.p cells in the same animal.

We then asked whether the duration of the BAR-1 pulse depended on the time of pulse onset. Because the duration of the L2 larval stage varied significantly between strains and animals, we examined the pulse onset time *t*_0_*/T*_L2_and duration *T/T*_L2_ relative to the duration of the larval stage, *T*_L2_. In this case, we found a striking anti-correlation, with late pulse onset resulting in shorter pulses (Fig. 5f). In fact, the data for all strains clustered along the line *T/T*_L2_ = 0.66 − *t*_0_*/T*_L2_, meaning that the end of the BAR-1 accumulation occurs at 66% of the L2 larval stage independent of the BAR-1 pulse onset time. This correlation also held for the *bar-1(*++*);cwn-1(0)* mutant, where not only the onset of the BAR-1 pulse was delayed but also the L2 larval stage was much extended in some animals. Overall, this indicated that the BAR-1 pulse ended at specific point in the L2 larval stage, with much less variability in timing compared to start of the pulse.

### Bias of cell fate decision by absolute BAR-1/β-catenin level

The observation of BAR-1 pulse dynamics raised the question how this controlled cell fate outcome. It is often assumed that the outcome of Wnt signaling depends on the absolute β-catenin level detected by the cell (Sawa and Korswagen 2013). In this case, both changes to BAR-1 pulse slope and pulse onset could bias hyp7/fusion frequency, by modulating absolute BAR-1 levels at the time the decision to fuse or not is made. However, it has also been suggested that cells respond to fold change in rather than absolute level of β-catenin (Goentoro and Kirschner 2009). In this case, cell fate outcome would depend only on BAR-1 pulse slope. Finally, cells could respond to more complex features of BAR-1 pulse dynamics.

To address this question, we first asked whether the change in hyp7/fusion frequency we observed between different strains was accompanied by systematic changes in pulse dynamics. Indeed, while *bar-1(*++*);cwn-1(0)* animals exhibited the same slope distribution as *bar-1(*++) animals (Fig. 5c), their BAR-1 pulse started at a later point in larval stage compared to *bar-1(*++) and *bar-1(*++*);lin-39(lf)* animals (Fig. 5d). This already suggested that the onset of the BAR-1 pulse was delayed compared to the time of fusion in this strain, but left open the possibility that the time of fusion was also delayed. Therefore, we attempted to directly measure the delay between pulse onset and time of fusion in these strains.

Because cells that fused did so rapidly after the onset of the BAR-1 pulse, it proved challenging to accurately determine the pulse onset time for these cells from our fitting procedure (see Methods). To circumvent this issue, we selected animals where one Pn.p cell, either P3.p or P4.p, assumed hyp7/fusion fate whereas the other assumed VPC fate, in *bar-1(*++*);lin-39(lf)* and *bar-1(*++*);cwn-1(0)* animals where both P3.p and P4.p frequently assumed hyp7/fusion fate. We then compared within the same animal the fusion time *t*_fusion_ in the hyp7/fusion cell with the pulse onset time *t*_0_ in the VPC (Fig. 6a). For *bar-1(*++*);lin-39(lf)* animals, the time of fusion correlated strongly with the pulse onset time (R=0.86 for P3.p assuming hyp7/fusion and P4.p VPC fate). Specifically, the data clustered along the line *t*_fusion_*T*_L2_ = 0.2 + *t*_0_/*T*_L2_, i.e. cell fusion occurred at a time 0.2*T*_L2_, or on average ∼2 hrs, after the onset of the BAR-1 pulse. However, in *bar-1(*++*);cwn-1(0)* animals, where the hyp7/fusion frequency is increased, the delay was halved, with cell fusion now occurring at a time 0.1*T*_L2_after onset of the BAR-1 pulse (Fig. 6a). Because *bar-1(*++*);cwn-1(0)* animals generally have similar BAR-1 pulse slopes as the *bar-1(*++) and *bar-1(*++*);lin-39(lf)* mutants (Fig. 5c), the main impact of the observed shorter delay between pulse onset and time of fusion would be to lower inhibitory BAR-1 levels at the time cell fusion is activated. In agreement with this, when we compared BAR-1 levels, as measured by BAR-1::GFP fluorescence, between *bar-1(*++*);cwn-1(0)* and *bar-1(*++*);lin-39(lf)* animals, we found lower BAR-1 levels in fusing P3.p and P4.p cells in the *cwn-1(0)* background at the time of cell fusion (Fig. 6b). Indeed, this observed difference is predicted when absolute BAR-1 and LIN-39 levels bias cell fate outcome in an additive manner: because the fusion inhibitor LIN-39 is not functional in *bar-1(*++*);lin-39(lf)* mutants, higher BAR-1 levels would be required to inhibit fusion.

**Figure 6.**
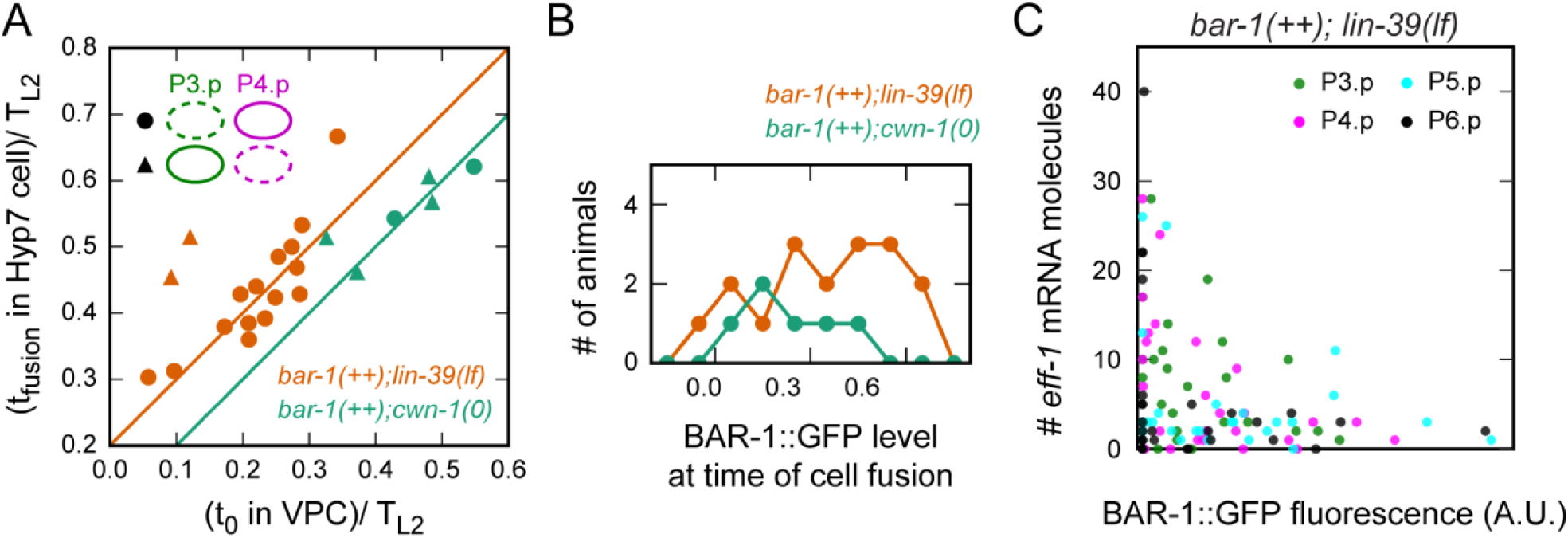
Impact of BAR-1 dynamics on cell fate decision. **(A)** Overview of interactions. Expression of *eff-1* and cell fusion are induced through an unknown activating signal, while BAR-1/β-catenin and LIN-39/Hox inhibit *eff-1* expression by inducing inhibitors, such as EGL-18 and ELT-6 (Koh et al. 2002), Developmental timing cues control both activation of *eff-1* expression in the mid-L2 stage and BAR-1 accumulation dynamics, with the BAR-1 accumulation pulse starting and ending in the early and late L2 stage, respectively. **(B), (C)** Examples linking the experimentally observed variability in LIN-39 levels and BAR-1 pulse dynamics to cell fate outcome. If combined levels of fusion inhibitors LIN-39 (green line) and BAR-1 (red line) are sufficiently high at the time of *eff-1* activation (blue line), no cell fusion occurs (B). Reduced levels of LIN-39 and BAR-1 allow for activation of *eff-1* expression, resulting in cell fusion (C). BAR-1 levels can be reduced both by delayed BAR-1 pulse onset time for a constant BAR-1 accumulation rate (C, top panel), or by lowering the BAR-1 pulse slope, corresponding to lower BAR accumulation rate (C, bottom panel).

Finally, we examined the relation between absolute BAR-1 levels and *eff-1* expression. In general, we expected *eff-1* expression to depend both on BAR-1 and LIN-39 level. To avoid confounding the relation between BAR-1 level and *eff-1* expression by variability in LIN-39 level, we quantified *eff-1* mRNA level in *bar-1(*++*);lin-39(lf)* animals. In general, we found that unfused P(3-6).p cells with high BAR-1::GFP level had few *eff-1* transcripts, while high *eff-1* levels were found only in unfused P(3-6).p cells with low BAR-1::GFP levels (Fig. 6c). Overall, these results are consistent with BAR-1 biasing the hyp7/fusion versus VPC fate decision through its absolute level.

## Discussion

### Variability in LIN-39/Hox and BAR-1/β-catenin dynamics

Here, we combined a novel time-lapse microscopy approach with quantitative analysis to study how the stochastic outcome of the hyp7/fusion versus VPC fate decision in the P3.p cell is controlled by variability in the dynamics of the underlying signaling network (Fig. 7). We found evidence that linked variability in the dynamics of the fusion inhibitors LIN-39/Hox and BAR-1/β-catenin to cell fate outcome. First, we observed that LIN-39::GFP levels exhibited significant variability, both in time and between animals (Fig. 3c,d). In addition, in *lin-39::gfp;cwn-1(0)* animals, where both P3.p and P4.p assume hyp7/fusion fate in a stochastic manner, we found that cell fusion occurred in the cell with the lowest LIN-39::GFP level (Fig. 3f). Second, we found that BAR-1/β-catenin accumulated in P(3-6).p cells in a single, ∼1-4hr pulse around the time of the decision between vulva precursor or hyp7/fusion fate (Fig. 4). These BAR-1 pulses showed considerable animal-to-animal variability, both in time of onset and in slope, i.e. the rate of BAR-1 accumulation (Fig. 5a-d). BAR-1 pulses likely impacted cell fate outcome by modulating the absolute BAR-1 level at the time of activation of cell fusion. In particular, two observations suggested that BAR-1 impacts fusion frequency through its absolute level: Pn.p cells in a mutant lacking the parallel fusion inhibitor LIN-39 fused with higher BAR-1::GFP levels at time of cell fusion, compared to a strain where LIN-39 was present (Fig. 6b). Moreover, high expression of *eff-1*, the fusogen protein required for cell fusion, only occurred in Pn.p cells with low BAR-1::GFP levels (Fig. 6c).

**Figure 6.**
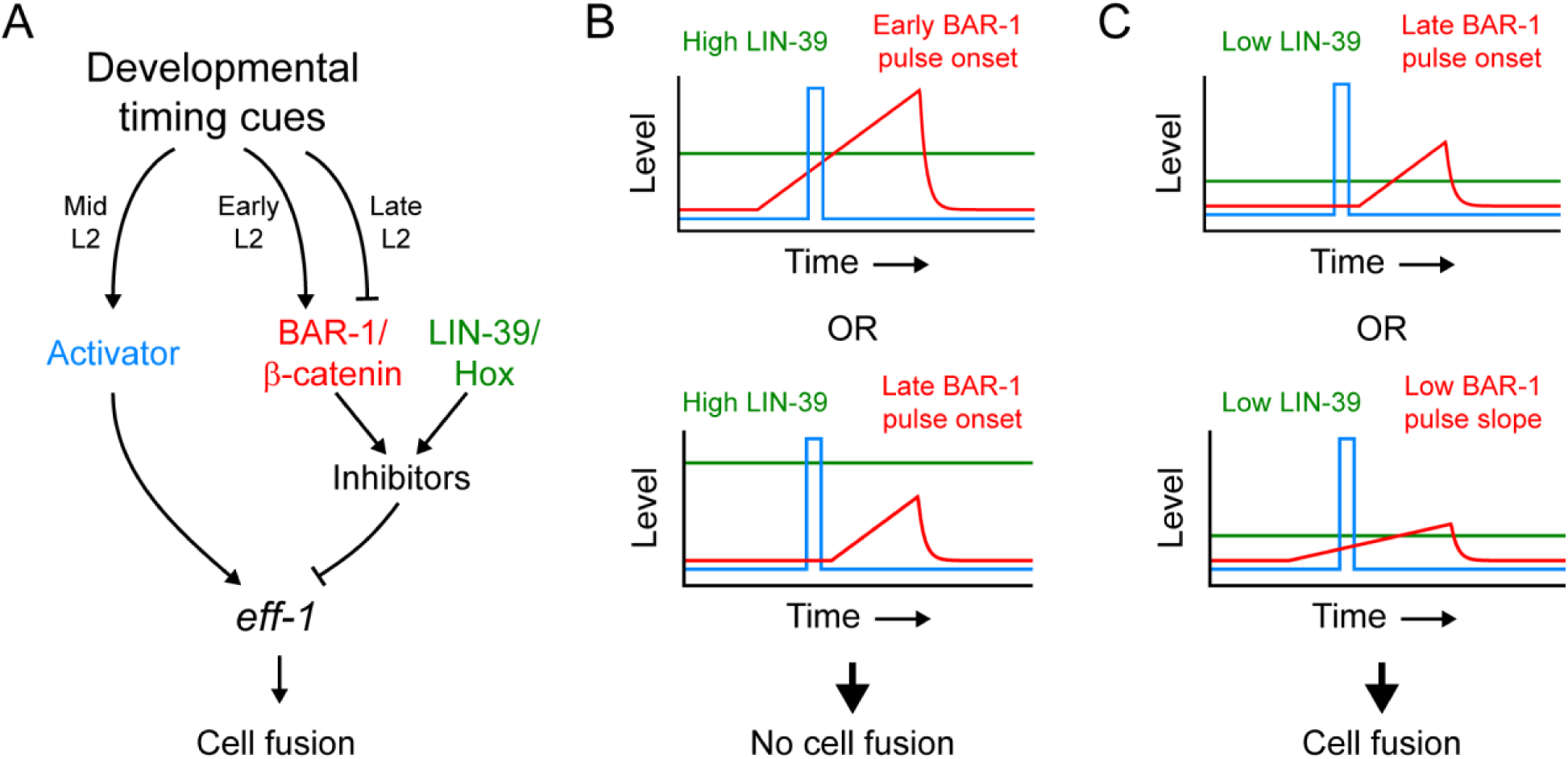
Model of cell fate bias by variability in LIN-39 levels and BAR-1 pulse dynamics. **(A)** Correlation between pulse onset time *t*_0_ in a VPC and fusion time *t*_fusion_ in a hyp7/fusion cell. Circles correspond to animals where P3.p, but not P4.p, assumed hyp7/fusion fate and triangles to animals where the cell fate assignment is reversed. The lines correspond to *t*_fusion_ΔT + *t*_0_*T*_L2_, with Δ*T* = 0.2 (orange) and Δ*T* = 0.1 (green). The *bar-1(*++*);cwn-1(0)* mutant has a shorter time delay Δ*t* between BAR-1 pulse onset and fusion. **(B)** Comparing BAR-1 levels at the time of cell fusion between *bar-1(*++*);lin-39(lf)* (orange, n=17) and *bar-1(*++*);cwn-1(0)* animals (green, n=6). Data for fusing P3.p and P4.p cells was pooled. Even though the difference in the mean of the two distributions was not significant due to the limited number of fusion events that could be recorded (Independent t-test, P=0.2), the highest BAR-1::GFP levels at the time of cell fusion were observed in the *bar-1(*++*);lin-39(lf)* mutant that lacks the parallel fusion inhibitor LIN-39. This was consistent with LIN-39 and BAR-1 inhibiting cell fusion in an additive manner through their absolute levels. **(C)** Number of *eff-1* mRNA molecules, as measured using smFISH, and BAR-1::GFP fluorescence level in unfused P(3-6).p cells of *lin-39(lf);bar-1(*++) animals. Each marker corresponds to a single cell (n=28 total animals). High *eff-1* expression (>20 mRNA molecules) was only observed for sufficiently low BAR-1::GFP levels.

### Temporal regulation

Our observations revealed a high degree of temporal regulation of the key events controlling the decision to fuse or not (Fig. 7a). We found that mutants that strongly impacted hyp7/fusion frequency had little impact on the time of cell fusion (Fig. 1e) and cell fusion occurred in a synchronized manner in mutants where multiple Pn.p cells assume hyp7/fusion fate (Fig. 1f). Similarly, we found that BAR-1 pulses started and ended at distinct points in the L2 larval stage, with their timing synchronized between Pn.p cells (Figs. 4, 5e). The onset and end of BAR-1 pulses showed clear differences in terms of variability: whereas BAR-1 pulse onset varied strongly between animals, BAR-1 pulses ended at a much more specific point in the L2 larval stage, leading to a broad distribution of BAR-1 pulse durations (Fig. 5e,f). Our findings linked the timing of BAR-1 pulses to the frequency of cell fusion. In particular, the only detectable difference between a *bar-1(*++) mutant, with low fusion frequency, and a *bar-1(*++*);cwn-1(0)* mutant, with high fusion frequency, was a delay in BAR-1 pulse onset with respect to the time of cell fusion (Figs. 5c,d, 6a). The strong link between the timing of cell fusion and BAR-1 pulse dynamics with L2 larval stage progression suggests that these events are regulated by a timing program that is independent of LIN-39 and Wnt signaling, either directly by a body-wide timing program such as the heterochronic pathway (Resnick, McCulloch, and Rougvie 2010) or indirectly, e.g. as function of Pn.p cell cycle progression during the L2 larval stage.

### Control of *eff-1* expression

LIN-39 and Wnt signaling likely influence the decision to assume hyp7/fusion fate by controlling expression of the fusogen protein EFF-1 (Fig. 7a). Indeed, we found that *eff-1* expression was generally low (< 10 mRNA molecules per cell), but showed a strong peak (20-60 mRNA molecules per cell) shortly before fusion (Fig. 2). Moreover, LIN-39 and Wnt signaling mutations that strongly impacted fusion frequency did not lead to an overall increase in *eff-1* mRNA level, but rather changed the fraction of animals in which we observed an *eff-1* expression peak (Fig. 2d). The change in *eff-1* expression in LIN-39 and Wnt mutants was consistent with their known function as inhibitors of cell fusion (Gidi Shemer and Podbilewicz 2002; D M Eisenmann et al. 1998). However, because LIN-39 and BAR-1 are typically assumed to act as activators, their inhibitory action is probably mediated by the induction of transcription factors that directly inhibit *eff-1* expression. For LIN-39, this likely occurs through the transcription factors EGL-18 and ELT-6 (Koh et al. 2002), but whether BAR-1 impacts *eff-1* expression via these same factors remains an open question. Even though we found evidence linking (variability in) both LIN-39 and BAR-1 levels to cell fate outcome, these results were obtained in strains where LIN-39 and BAR-1 were overexpressed. As a consequence, it remains an open question how strong their relative contributions to the bias of hyp7/fusion versus vulva precursor cell fate are under wild-type conditions. Finally, no activator of *eff-1* expression is currently known. However, our observation that timing of cell fusion is not impact by loss of LIN-39 or Wnt signaling (Fig. 1e) implied the presence of one or more unknown activators that induce expression of *eff-1* at a highly specific point in the L2 larval stage. Overall, our observations suggested that hyp7/fusion fate required sufficiently high *eff-1* expression and that a mixture of activating and inhibitory signals controlled the fraction of animals in which this threshold was crossed and cell fusion occurred.

### Model of stochastic cell fate bias by LIN-39/Hox and BAR-1/β-catenin

Based on the above observations we currently favor the following model for how variability in LIN-39 and BAR-1 dynamics biases the stochastic decision to assume hyp7/fusion or vulva precursor fate (Fig. 7b,c): *eff-1* expression is induced by an activating signal at a highly specific time, but simultaneously inhibited by LIN-39 and BAR-1 in a manner proportional to their absolute levels in the cell. If the combined level of LIN-39 and BAR-1 is sufficiently high, *eff-1* expression remains below threshold required for fusion and the cell assumes vulva precursor cell fate (Fig. 7b). However, the observed variability in LIN-39 and BAR-1 dynamics can cause their inhibitory levels to fall below what is required to block *eff-1* expression (Fig. 7c). For BAR-1 pulses, this could occur not only by a lower rate of BAR-1 accumulation, i.e. low BAR-1 pulse slope, but also by delaying the onset of the BAR pulse relative to the time of activation of *eff-1* expression. Hence, in this model it is not the absolute BAR-1 level itself, as is generally assumed, but rather the timing of its accumulation dynamics relative to other developmental events, that is a key factor determining the stochastic cell fate outcome.

### Generation of BAR-1/β-catenin pulses

The observation that BAR-1 accumulated in Pn.p cells in precisely timed pulses was surprising. However, pulsatile β-catenin accumulation dynamics under constant external Wnt levels has been observed in recent years (Murphy et al. 2014; Kafri et al. 2016; Massey et al. 2019). An important question is whether the BAR-1 pulse dynamics we observed here is influenced by the elevated levels of BAR-1 in the integrated transgene in which it was visualized. While our endogenously tagged BAR-1::GFP line had wild-type rates of hyp7/fusion fate in P3.p cells, we were unable to detect GFP fluorescence in Pn.p cells using our time-lapse microscope, indicating significantly lower expression levels compared to the BAR-1::GFP transgene. In the canonical Wnt signaling pathway, BAR-1 levels are controlled by their degradation through the β-catenin destruction complex, with Wnt signaling leading to inactivation of this complex and accumulation of BAR-1 (Sawa and Korswagen 2013). In the overexpression BAR-1::GFP strain, the absence of BAR-1::GFP prior to the pulse and its rapid disappearance directly afterwards showed that the increased BAR-1 level was still sufficiently low to not overwhelm the β-catenin destruction complex. Hence, if the BAR-1 accumulation pulse is controlled by the activity of the destruction complex, as is expected, BAR-1 should exhibit the same pulse dynamics under wild-type conditions.

It is an important question how the BAR-1/β-catenin pulse is generated. BAR-1/β-catenin accumulation could be controlled by changes in the level of the Wnt ligands outside of the cell or rather by changes in the Wnt pathway components inside the cell, such as changes to Wnt receptor levels or presence/activity of components of the β-catenin destruction complex. We currently favor the latter hypothesis, since neuronal cells close to the Pn.p cells show BAR-1::GFP expression in the late L1/early L2 stage when Pn.p cells do not, with a significantly decreased BAR-1::GFP signal in those cells in a *cwn-1(0)* background (data not shown). This suggests that Wnt ligands are already present and able to activate Wnt signaling at this time and position in the body.

As ligand-activated Wnt receptors sequester and thereby inactivate the destruction complex that induces β-catenin degradation, we expected that changing Wnt levels would predominantly impact the rate of β-catenin accumulation (Sawa and Korswagen 2013; Clevers and Nusse 2012). However, using the *cwn-1(0)* mutant we found that removing a Wnt ligand instead changed only the timing of the induced BAR-1/β-catenin pulse (Figs. 4d,f and 5c,d), which is difficult to explain based on our current knowledge of the Wnt pathway. Particularly surprising is that loss of *cwn-1* almost completely removed the variability in BAR-1/β-catenin pulse timing between adjacent Pn.p cells in the same animal (Fig. 5e). In absence of CWN-1, other Wnt ligands such as EGL-20 repress hypodermal fate, albeit at reduced efficiency (Pénigault and Félix 2011b). Our results indicate that CWN-1 acts in a significantly more stochastic manner, either on the level of ligand/receptor interaction or the delivery of ligands to the Pn.p cells, than the other Wnt ligands in the body, even though it is the Wnt ligand expressed closest to the Pn.p cells (David M Eisenmann 2005; Harterink et al. 2011).

### Role of β-catenin pulse dynamics in development

It is increasingly clear that many of the canonical metazoan signaling pathways control development using temporal information encoded in their dynamics (Levine, Lin, and Elowitz 2013; Shimojo, Ohtsuka, and Kageyama 2008) In particular, pulses in the output of signaling pathways have now been identified *in vivo* in a number of developmental systems. For example, EGF signaling acts in a pulsatile manner in the VPCs, with signaling strength transmitted in the frequency of pulses instead of a continuous graded signal (de la Cova et al. 2017). Moreover, time-lapse imaging of oscillatory, rather than pulsatile dynamics of Notch and Wnt signaling during segmentation of mouse embryos showed that the relative phase of the two oscillations instructs the segmentation process (Sonnen et al. 2018). Although single pulses of β-catenin accumulation have been observed in cell lines following exogenous application of Wnt ligands (Murphy et al. 2014; Kafri et al. 2016), their function inside in context of development is more poorly understood. Hence, the BAR-1 pulses we observed here during C. elegans larval development could be an attractive model to study how pulsatile Wnt signaling dynamics impacts cell fate decisions.

It remains an open question what the advantage is of activating BAR-1/β-catenin as a pulse rather than it being present continuously like the parallel hypodermal fate inhibitor LIN-39. One potential advantage is that BAR-1/β-catenin pulsatile dynamics allows for pulse timing as an additional control parameter, next to BAR-1/β-catenin accumulation rate, to tune cell fate frequency (Fig. 7b,c). Whether cells receive Wnt input is tightly controlled in space, e.g. by regulating Wnt receptor expression. Pulsatile dynamics could be a powerful mechanism to control precisely when cells respond to ligands in time. This might be particularly important because the same signaling pathways are used many times during development, sometimes even in the same cell, to control different developmental events. Reading out these signaling pathways only at particular time points would allow the reconfiguration of the pathway from executing one developmental decision to another. Interestingly, Wnt signaling is used in VPCs at the mid-L3 stage, ∼10 hrs after the hypodermal versus VPC decision, to control the anteroposterior orientation of their asymmetric divisions (Green, Inoue, and Sternberg 2008). Here, EGL-20 plays an important role, whereas BAR-1 and CWN-1 have a smaller contribution. The decay of the BAR-1/β-catenin pulse at the end of the L2 stage might be crucial to avoid temporal crosstalk between the outputs of the Wnt pathway as the VPCs transition from one process to the next.

In conclusion, we have shown here that β-catenin accumulation can be highly dynamic during development, with temporal information instructing development contained in its dynamics. Many (stochastic) cell fate decisions in organism from nematodes to humans are controlled by Wnt signaling and it will be interesting to see whether pulsatile β-catenin plays a similar role in biasing cell fate frequencies in those systems. The quantitative approach we employed here, combining in vivo time-lapse imaging of β-catenin dynamics with measurements of key dynamical parameters such as pulse slope and pulse timing, can provide a template for such future studies.

## Materials and Methods

### CONTACT FOR REAGENT AND RESOURCE SHARING

Further information and requests for resources and reagents should be directed to and will be fulfilled by the Lead Contact, Jeroen van Zon.

### EXPERIMENTAL DETAILS

#### Strains

All strains were handled according to the standard protocol on Nematode growth medium (NGM) agar plates with OP50 bacteria (Brenner 1974). Experiments were performed on L2 stage hermaphrodites. Strains were obtained from the CGC unless otherwise indicated. The following mutations were used in this study:

LGII: *cwn-1(ok546)* (The C. elegans Deletion Mutant Consortium 2012).

LGIII: *lin-39(gk893)* (The C. elegans Deletion Mutant Consortium 2012), *lin-39(n709)* (Clark, Chisholm, and Horvitz 1993).

LGX: *bar-1(ga80)* (D M Eisenmann et al. 1998).

The following transgenes were used in this study:

*ncIs13[ajm-1::GFP]* (Liu et al. 2005).

*sIs11337[rCesY37A1B.5::GFP* + *pCeh361)]* (McKay et al. 2003).

*ouIs20 [ajm-1::mCherry* + *unc-119*+*] (*gift from Alison Woollard).

*itIs37[pie-1p::mCherry::H2B::pie-1 3’UTR* + *unc-119(*+*)] IV, stIs10116[his-72(promoter)::his-24::mCherry::let-858 3’UTR* + *unc-119(*+*)], stIs10311[lin-39::TGF(3D3)::GFP::TY1::3xFLAG]*

(Sarov et al. 2012),

*gaIs45[pDE218(bar-1::bar-1::GFP)]* (gift from David Eisenmann) (D M Eisenmann et al. 1998),

*stIs10226[his-72p::HIS-24::mCherry::let-858 3’ UTR* + *unc-119(*+)] (Sarov et al. 2012).

The presence of the *cwn-1(ok546)* homozygous deletion was confirmed by nested PCR screening. The following primers were used: outer F (‘5-TCGTTTCTGACATGGCTCAC-3’), outer R (‘5-ACCCATCCTTTCCCAATCTC-3’), inner F (‘5-CGTATCCACGACCACAACAG-3’) and inner R (5’-AGAATCTTCACACCAACGGG-3’).

#### Time-lapse imaging

The microchamber size used in the study was 190 μm x 190 μm, with a depth of 10 μm, made as previously described (Gritti et al. 2016). Using an eyelash attached to a glass pipette, OP50 bacteria were used as “glue” to transfer eggs into the microchambers using M9 solution to keep the microchamber moist. A Nikon Ti-E inverted microscope with a large chip camera (Hamamatsu sCMOS Orca v2) and a 60 X magnification objective (NA=1.4 oil immersion) was used for imaging. Transmission imaging was performed using an LED light source (CoolLED pE-100 615nm), while 488 and 561 nm fluorescence images were acquired using Coherent OBIS LS 488-100 and OBIS LS 561-100 lasers, respectively. Images were acquired in a temperature controlled room at 19° with a sample temperature of 22°. Exposure time was 10 ms and approximately 25 images were taken with a z-distance of 1 μm. Images were taken every 20 min. Images were 2048 × 2048 pixels and saved in 16-bit TIFF format. Fusion times were determined by *ajm-1::GFP*/*mCherry* localization and morphology.

#### Quantitative Image Analysis

For all experiments, transmitted light images where used to identify molt times. Custom Python scripts (available at Github: https://github.com/jvzonlab/P3p_bar1) and ImageJ were used to quantitatively analyze the acquired images (Schindelin et al. 2012; Schneider, Rasband, and Eliceiri 2012). First, images to be used for quantitative analysis were corrected for uneven laser illumination in the following manner: flat field intensity for the particular experiment was obtained by imaging a uniformly fluorescent (488nm) testing slide, and averaging the result of 10 images. We divided each pixel’s intensity value in the experimental images by the corresponding flat field pixel’s intensity, normalized to the median value of the entire flat field image. This normalization procedure corrects for position-dependent variation in light intensity due to the Gaussian profile of the laser beam. The region of interest was cropped at this time. Pn.p cells were manually identified by stereotyped nuclear position location and the domains of *ajm-1::GFP/mCherry* expression, if present. To measure LIN-39::GFP expression, a mask was manually drawn around the nucleus and the mean fluorescence intensity of the pixels within the mask was calculated. The z-slice closest to the center of the nucleus was used. A background fluorescence measurement for each image was obtained by creating a mask of the intranuclear space in a region near P3.p and P4.p along the axis of the ventral nerve cord. The mean background fluorescence value was then subtracted from the mean fluorescence value of the reporter for the same image. To measure BAR-1::GFP expression, a mask was manually drawn around the Pn.p cytoplasmic region using AJM-1::mCherry signal as a positional guide, with background corrections performed similarly as described above. For the Supplementary Movies, fluorescence images were computationally straightened and aligned, using the animal’s body shape and position of the Pn.p cells as measured from the fluorescent markers.

#### Single-molecule fluorescence in situ hybridization (smFISH)

Probe design and smFISH hybridization to visualize *eff-1* mRNA transcripts were performed as previously described (Huelsz-Prince and van Zon 2017; Raj et al. 2008). Custom probes were designed against the exons of the *eff-1* gene by utilizing the Stellaris® RNA FISH Probe Designer (Biosearch Technologies, Inc., Petaluma, CA). The probes were hybridized with the dye Cy5 (Huelsz-Prince and van Zon 2017). The sequences of the oligonucleotide probes used in this study are listed in the Supplementary Methods. Animals were collected by washing plates with M9 and were fixed in 4% formaldehyde in 1 X PBS for 45 min at room temperature. Fixed animals were permeabilized in 70% ethanol at least one night at 4°C. Subsequently, animals were incubated with the 0.5 μl probes overnight at 30°C in Stellaris® Hybridization Solution containing 10% formamide. The next day, animals were quickly washed two times with 10% formamide and 2 X SSC, followed by an incubation wash for 30 min at 30°C. DAPI was added at 0.01 μg/ml in a final incubation step for 20 min at 30°C. Animals were mounted in Glox Buffer with catalase and glucose oxidase, and images were acquired with a Nikon Ti-E inverted fluorescence microscope, equipped with a 100X plan-apochromat oil-immersion objective and an Andor Ikon-M CCD camera controlled by μManager software (Edelstein et al. 2014). Stacks of each animal were taken with a z-distance of 0.33 μm and approximately 30 images were taken per animal. Cy5 exposure time was 3 s, while DAPI and GFP exposure time were 100 ms and 500 ms, respectively. Animals were then imaged at 40 X to determine their body length, which was measured using ImageJ by drawing a spline from the tip of the head to the end of the tail. smFISH images were analyzed with a custom Python script using techniques previously described (Raj et al. 2008). The *ajm-1::GFP* transgene was used to determine the cell fusion status. For Figure 6C, ImageJ was used to quantify GFP fluorescence in the Pn.p cellular region of P3-P6.p by selecting the slice most central to the nucleus. The cellular region was manually outlined and the mean fluorescence value of the cellular region was quantified using ImageJ. A background fluorescence measurement near the Pn.p cells for each image was then subtracted to correct for background fluorescence. If the corrected BAR-1::GFP fluorescence resulted in a negative value, it was manually adjusted to zero.

#### Parameterization and fitting of BAR-1::GFP dynamics

To fit the experimentally measured BAR-1::GFP dynamics, we assume the following minimal model of BAR-1 production and degradation:s

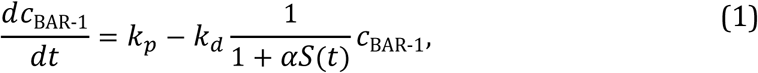

where *c*_BAR-1_ is the BAR-1 level and *k*_*p*_ is the BAR-1 production rate. In the absence of Wnt signaling, *S*(*t*) = 0, the degradation complex degrades BAR-1 at a basal rate *k*_*d*_ However, in the presence of Wnt signaling, *S*(*t*) = 1, the degradation complex is inhibited (David M Eisenmann 2005) and degradation occurs at a reduced rate 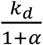, with *α* > 1. In the model, we assume the. BAR-1 pulse is generated by changes in Wnt signaling level *S*(*t*). In particular, we assume Wnt signaling is activated at a constant level *S*(*t*) = 1 starting at time *t*_0_ and ending at time *t*_0_ +*T*, where *T* is the pulse duration. For other times, Wnt signaling is not activated, *S*(*t*) = 0. These assumptions yield the following expression for the BAR-1 dynamics:

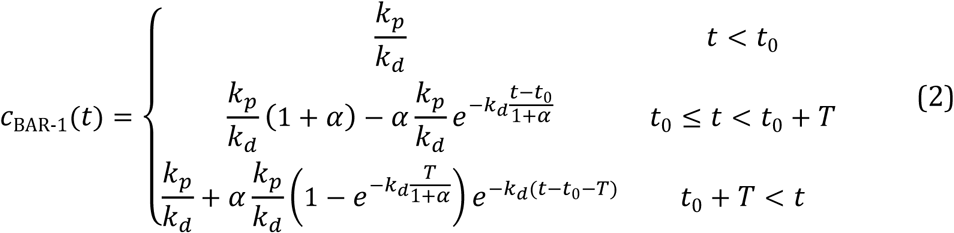

For sufficiently long pulse duration T, the BAR-1 level will reach a steady state 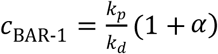. However, in the experimentally obtained data we never observed BAR-1::GFP levels reaching a steady state before the end of the pulse. Instead, we found that BAR-1::GFP accumulation remained approximately linear throughout the full duration of the pulse. Indeed, when the pulse duration is sufficiently short or the Wnt-mediated inhibition of BAR-1 degradation is sufficiently strong, i.e.(1 ʱ *α*) ≫ *k*_*d*_*T*, the exponential term in *c*_BAR-1_(t) for *t*_0_ ≤*t*<*t*_0_+*T* reduces to an expression that linearly with time in linear fashion, giving rise to the following expression used to fit the experimental data:

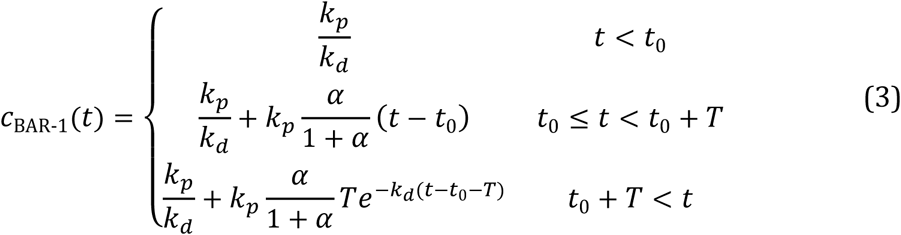

We fitted this expression to the experimental BAR-1::GFP data by least-square fitting, using the implementation of the Levenberg-Marquardt algorithm, as implemented in the Python function scipy.optimize.curve_fit (Eric Jones, Oliphant, and Peterson 2001) and using *k*_*p*_, *k*_*d*_, *α, t*_0_ and *T* as fit parameters. We obtained these fit parameters for each Pn.p cell by independent fitting. For fusing cells, we fitted *c*_BAR-1_(*t*) to the experimental data only for time points until the experimentally determined time of fusion, *t*_fusion_. We found that cell fusion always occurred before the end of the pulse, i.e. *t*_0_+*T*> *t*_fusion_ and, hence, was not defined for fusing cells. In general, this fitting procedure provides good fits for most BAR-1::GFP trajectories, but fails to converge to a correct fit for trajectories with very low pulse amplitude or no apparent pulse. In that case, we assume *c*_BAR-1_(*t*) = 0, with *t*_0_ and *T* not defined. To characterize pulse dynamics for non-fusing cell, we compare pulse onset time *t*_0_, pulse duration *T* and pulse slope 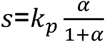. For fusing cells, pulse duration is not defined and instead we compare the time of fusion. Finally, it proved in practice to determine the time of pulse onset from the above fitting procedure in cells that fused. This was because fusion typically occurred only a few time points after the onset of the pulse, when BAR-1::GFP levels had not risen far above the background, providing too little data for the fitting procedure to converge on a proper fit. Using the observation that both pulse onset time and pulse slope were similar in neighboring Pn.p cells, we could still estimate the pulse onset tme in fusing cells by using the measured value from a neighboring non-fusing cell.

#### Generation of *bar-1::GFP* Using CRISPR-Cas9 Genome Editing

Wild type N2 *C. elegans* genetic background was used for the generation of the CRISPR/Cas9 *bar-1::GFP* allele. The endogenous *bar-1* gene was tagged C-terminally using the nested CRISPR technique (Vicencio et al. 2019). The protocol was followed with slight modifications. For step two of the nested CRISPR technique, instead of a fully double-stranded PCR product serving as the repair template, partially single-stranded dsDNA GFP donors were used. Briefly, two GFP PCR products were created (see Supplemental Methods for primers), one spanning the entire GFP coding region, while the other spanning the interior sequence. The PCR products were hybridized together to create a donor cocktail featuring fragments with ssDNA overhangs (56 and 57 bp) (Dokshin et al. 2018). The template for GFP was the plasmid pJJR82 (gift from Mike Boxem). The two GFP PCR products were annealed by mixing the individual PCR products 1:1, then running the following thermocycler program: 95° 2:00s, 85° 0:10s, 75° 0:10s, 65° 0:10s, 55° 1:00s, 45° 0:30s, 35° 0:10s, 25° 0:10s, and finally 4° hold. All crRNA (2 nmol product) and single-stranded repair templates (4 nmol product) were prepared commercially (IDT) and were resuspended in 20 or 40 µl, respectively, of nuclease-free duplex buffer to yield a stock concentration of 100 µM. The strand of the single-stranded repair template was determined by data from (Farboud, Severson, and Meyer 2019). Two crRNAs were used for step one of nested CRISPR, with the double-stranded breaks occurring 15 and 18 bp away from the insertion site. The first injection mix contained 250 ng/µl Cas9 (IDT), 10 ng/µl tracrRNA, 3 ng/µl *bar-1* crRNA #1, 3 ng/µl *bar-1* crRNA #2, 3 ng/µl GFP crRNA, 2 ng/µl *bar-1* repair ssODN, 400 ng/µl *GFP* PCR product, 1 ng/µl *dpy-10* crRNA, 1 ng/µl *dpy-10(cn64)* repair ssODN, and RNAse free buffer to a final volume of 10 µl. The second injection mix contained 250 ng/µl Cas9 (IDT), 10 ng/µl tracrRNA, 7 ng/µl GFP crRNA, 2 ng/µl *bar-1* repair ssODN, 400 ng/µl *GFP* hybrid PCR product, 3 ng/µl *dpy-10* crRNA, 1 ng/µl *dpy-10(cn64)* repair ssODN, RNAse free buffer to 10 µl. In both steps, young adult N2 hermaphrodites (P0) were injected in the germline using an inverted micro-injection setup (Eppendorf). After injection, *dpy-10* edited animals (F1) were singled and grown at 20°, and the *bar-1* locus was genotyped for successful co-conversion by PCR amplification with OneTaq polymerase (New England Biolabs) using primers that annealed to the genomic DNA surrounding the insertion point. Potential GFP insertions were determined visually by amplicon size on agarose gel and were sequenced to verify in-frame insertions (Macrogen Europe). The *dpy-10* edited co-CRISPR allele was removed from the genetic background from the first nested CRISPR step by picking a wild-type animal from a heterozygous *dpy-10* edited animal. In the second step, it was removed by outcrossing the edited *dpy-10* homozygous animal to N2 for one generation. Growth, movement, and fertility were all superficially wild-type in the tagged line.

## Data availability statement

Strains and plasmids are available upon request. Supplemental data (analyzed time-lapse and smFISH data for individual animals, fitted pulse parameter for BAR-1) and Python software used for analysis and creating figures are available at http://github.com/jvzonlab.

## Acknowledgements

This work is part of the research program of the Netherlands Organisation for Scientific Research (NWO) and was performed at the research institute AMOLF. Some *C. elegans* strains were provided by the CGC, which is funded by NIH Office of Research Infrastructure Programs (P40 OD010440). The work was supported by a European Research Council Starting Grant (338200-STOCHCELLFATE) awarded to J.S.v.Z.

## Author Contributions

Conceptualization, J.S.v.Z.; Methodology, J.R.K. and J.S.v.Z.; Formal Analysis: J.R.K., J.T, and J.S.v.Z.; Investigation, J.R.K., and J.T.; Writing – Original Draft, J.R.K. and J.S.v.Z.; Writing – Review & Editing, J.R.K. and J.S.v.Z.; Funding Acquisition, J.S.v.Z.; Supervision, J.S.v.Z.

---

**Figure S1.**
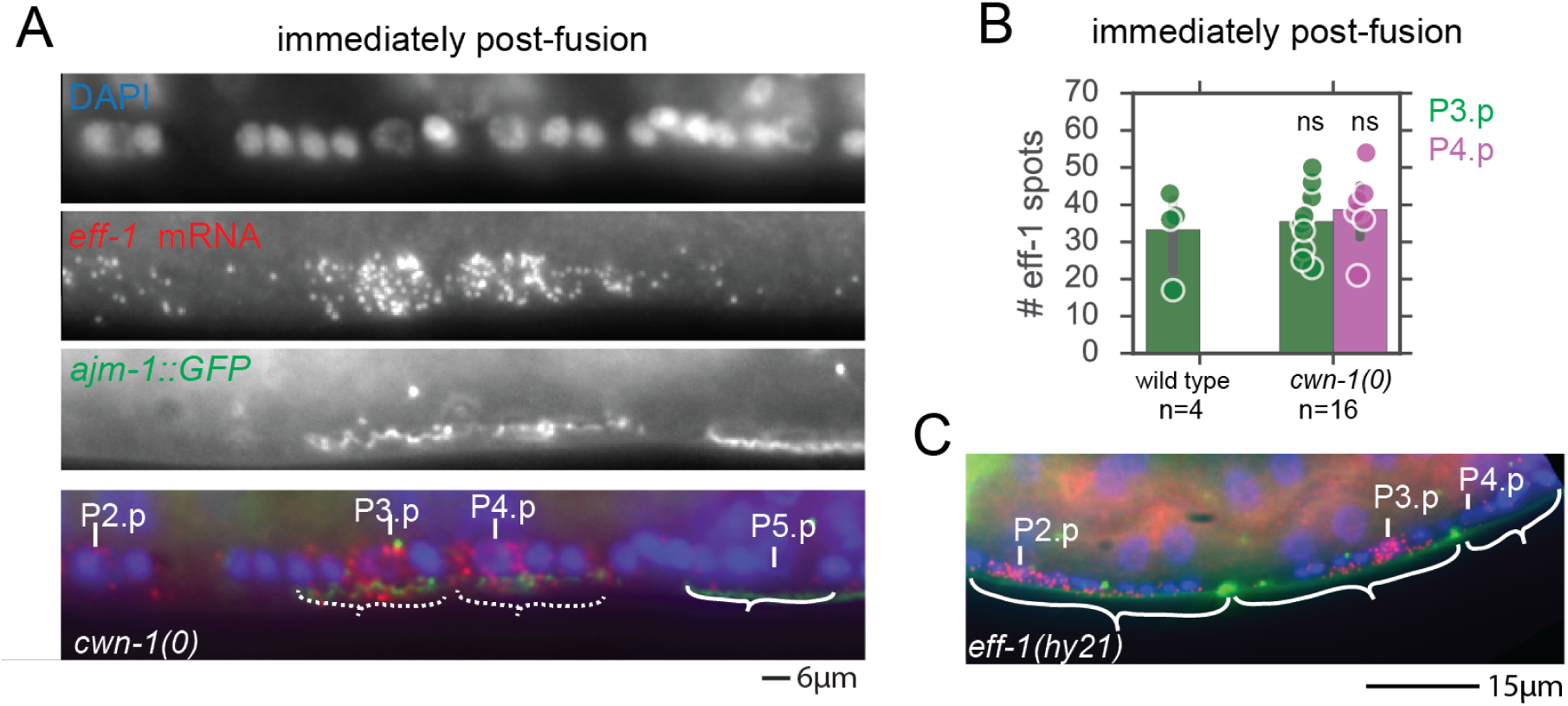
The gene *eff-1* is expressed at a high level before cell fusion. **(A)** Example of a *cwn-1(0)* animal with both P3.p and P4.p showing evidence of recent fusion, due to the currently degrading and ruffled appeared of AJM-1::GFP, marked by dashed-white brackets in the composite image (bottom panel). **(B)** Quantification of the number of *eff-1* spots in cells that have recently fused as in (A). Different genetic backgrounds show similar levels of *eff-1* expression at the time of fusion. Dots correspond to individual cells and are plotted with the mean and standard error. No significant differences are present in the average number of *eff-1* spots (Student’s t-test). **(C)** Image of a mid-L2 stage *eff-1(hy21)* temperature-sensitive-mutant with a point mutation that renders the protein non-functional, but allows for mRNA staining with smFISH probes. High levels of *eff-1* expression are seen in P2.p and P3.p, despite them remaining unfused (marked by white brackets).

## Supplemental Movies

**Movie S1**: (Top panel) Apical junction marker AJM-1::mCherry and (bottom panel) GFP expressed in the hypodermal syncytium hyp7 for a P3.p cell assuming hyp7/fusion fate. At the time of fusion with hyp7, 6h 20m after the start of the L2 larval stage, pronounced ruffling of AJM-1 is followed by its removal from P3.p. Concomitantly, GFP flows from the hyp7 syncytium into P3.p, as indicated by the yellow arrow. A *dpy-7p::mCherry* nuclear marker was used for cell identification (top panel).

**Movie S2**: (Top panel) Apical junction marker AJM-1::mCherry and (bottom panel) GFP expressed in the hypodermal syncytium hyp7 for a P3.p cell that assumes vulva precursor cell fate. AJM-1 is present in P3.p throughout the entire L2 larval stage and no inflow of GFP into P3.p is observed. A *dpy-7p::mCherry* nuclear marker was used for cell identification (top panel).

**Movie S3**: Pulsatile BAR-1 dynamics in P(3-5).p cells in a single animal. Time is relative to the start of the L2 larval stage. Each panel corresponds to a single cell. Shown are the apical junction marker AJM-1 (magenta) and BAR-1::GFP (green). The animal examined corresponds to that shown in Fig. 4a,b.

### DNA sequences used for CRISPR/Cas9 genome editing

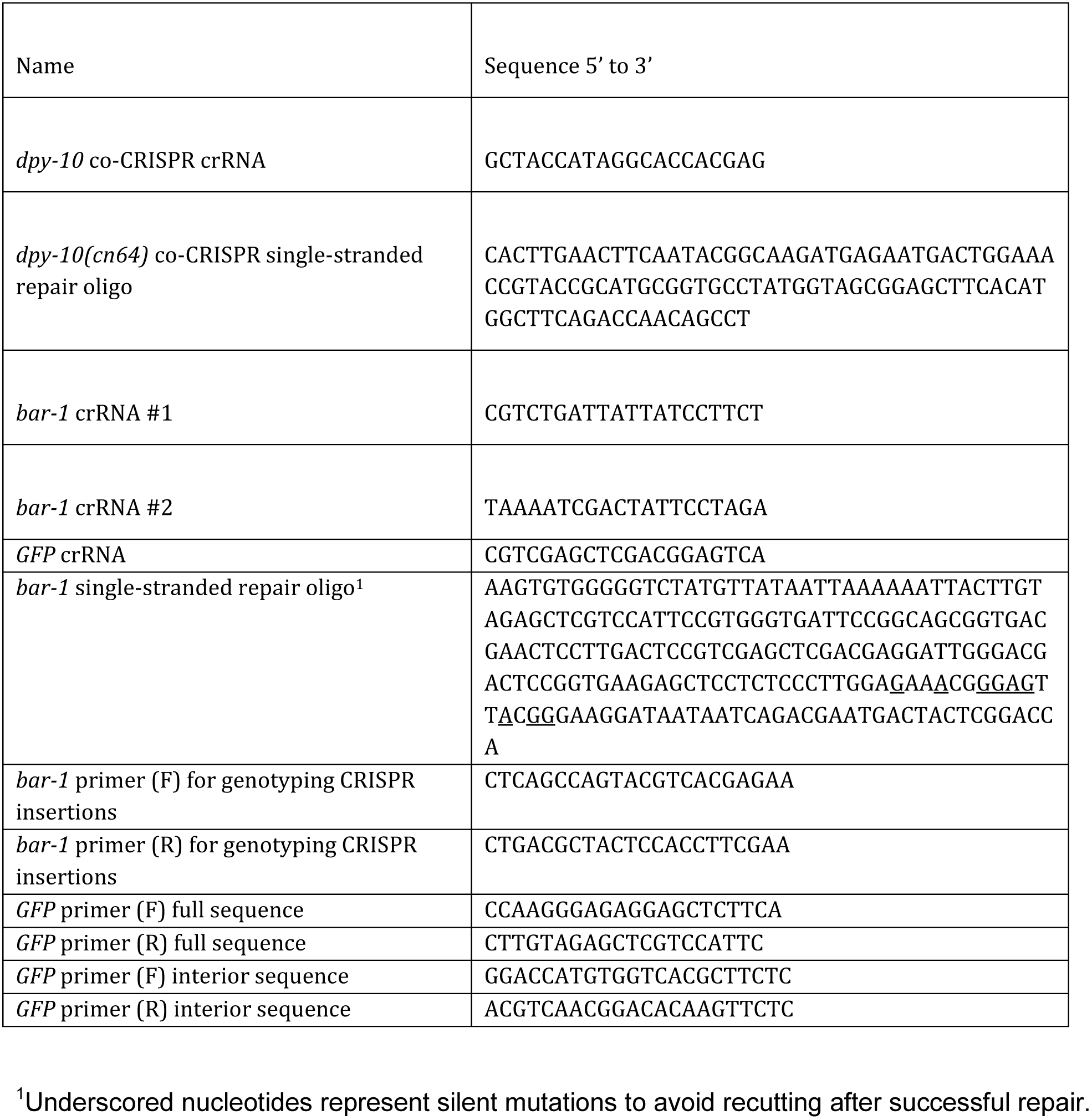

### smFISH probes used to hybridize with *eff-1* mRNA molecules

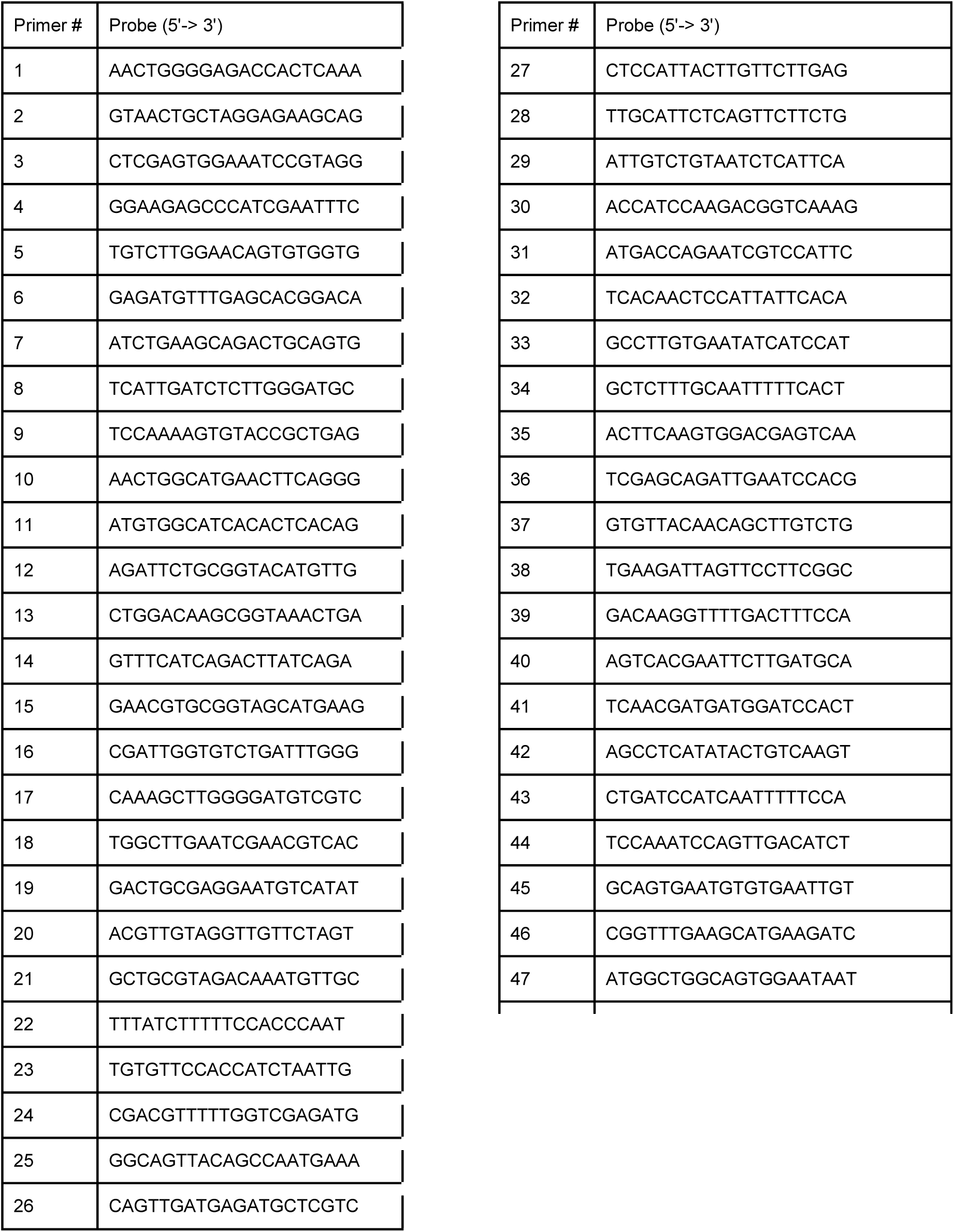

